# Knock-in ≠ knock-out: differential fitness effects of *cardinal* mutations in *Anopheles stephensi*

**DOI:** 10.64898/2026.07.07.737011

**Authors:** Mireia Larrosa-Godall, Lewis Shackleford, Philip T. Leftwich, Estela Gonzalez, Joshua X. D. Ang, Matthew Edwards, Katherine Nevard, James C.Y. Luk, Morgan Mckee, Rob Noad, Michelle A. E. Anderson, Luke Alphey

## Abstract

The kynurenine pathway metabolizes tryptophan into 3-hydroxykynurenine (3-HK), a precursor for ommochrome eye pigments synthesized via the *cardinal* (*cd*) gene in mosquitoes. While *cd* disruption was presumed neutral, we observed fitness costs in *Anopheles stephensi* knock-in but not knock-out *cd* mutants. Here we investigated this anomaly further by assessing survival, fecundity, and midgut integrity across multiple *cd* mutant lines. Heterozygous knock-in lines, expressing a fluorescent marker and guide RNA for CRISPR/Cas9, exhibited reduced survival post-blood feeding, larva-to-adult survival deficits, and midgut barrier dysfunction, whereas knock-outs showed no such costs. Oral supplementation with xanthurenic acid partially rescued knock-in mortality, implicating oxidative stress linked to 3-HK metabolism. Expression analyses suggest transgene insertion effects, rather than *cd* disruption, underlie these fitness costs. These findings highlight the importance of evaluating insertional effects in gene drive target selection and support *cd* as a viable target for genetic control strategies in *An. stephensi*.

## Introduction

The kynurenine pathway is a key metabolic route in mosquitoes, responsible for breaking down tryptophan (Trp) into several biologically active metabolites including 3-hydroxykynurenine (3-HK) (1). In this pathway, Trp is initially converted to N-formylkynurenine by the enzyme tryptophan-2,3-dioxygenase (TDO). N-formylkynurenine is then converted to kynurenine (Kyn) either spontaneously or via kynurenine formamidase (KFase), and subsequently to 3-HK by kynurenine monooxygenase (*kmo*, also known as kynurenine hydroxylase (*kh*)). 3-HK is readily oxidised under physiological conditions (1), leading to the formation of free radicals which have been linked to retinal damage (2), death of midgut epithelial cells (3), neurotoxic and neurobehavioral effects (4) and reduced lifespan (5) in invertebrates. Similar harmful effects have been observed in vertebrates, where accumulation of 3-HK is associated with aging and neurodegenerative disorders (6). Thus, tight regulation of 3-HK levels is crucial for organismal health.

In vertebrates, 3-HK is mainly hydrolysed to 3-hydroxyanthranilic acid (3-HAA) by kynureninase (1) or converted to xanthurenic acid (XA) by kynurenine aminotransferase (7). In contrast, mosquitoes lack kynureninase; instead 3-HK is converted to XA by 3-hydroxykynurenine transaminase (HKT) (1). XA is of particular interest in Anopheline mosquitoes as it has been reported to induce exflagellation of *Plasmodium* microgametocytes (8), and HKT has been identified as a potential target for vector control due to its role in larval survival (9). 3-HK also serves as the initial precursor for the ommochrome pathway, in which ommochrome pigments — the major eye pigment in mosquitoes (10) — are produced via nonenzymatic auto-oxidation (11,12) or by phenoxazinone synthase (PHS), encoded by the *cd* gene first identified in *Drosophila* (13). Mutations in this pathway generate distinctive eye-colour phenotypes, making these enzymes and substrates useful genetic markers for developing genetic transformation technologies in insects (14–18).

Recent advances in genetic manipulation, particularly the CRISPR–Cas9 system, have opened new avenues for controlling mosquito populations and reducing disease transmission (19,20). Cas9, guided by a sequence-specific single guide RNA (sgRNA), generates targeted double-stranded breaks in DNA that are repaired by either homology-directed repair (HDR) or the more error-prone end joining (EJ). Homing gene drives exploit HDR to bias inheritance, enabling the spread of a desired trait through a population at rates exceeding those expected under Mendelian inheritance, even if the modification imposes a fitness cost. These systems are designed for population suppression (reducing or eliminating mosquito populations) or for population modification (introducing traits that make mosquitoes resistant to pathogen infection or transmission). The success of such systems depends on identifying suitable genetic targets that can be efficiently edited without imposing severe fitness costs.

Genes involved in the ommochrome pathway have been considered broadly neutral, with disruption affecting the eye colour phenotype, but not impairing fertility or survival. This neutrality makes them attractive targets for Cas9-based gene drives, as mutations can be easily detected visually, for example mosaic eye-colour phenotypes resulting from somatic Cas9 activity (16,18). However, recent studies have challenged this assumption; introducing a gene drive into *kmo* in *An. stephensi* and generating a knock-out in *Ae. aegypti* led to reduced survival after blood feeding and decreased reproduction in homozygous females (1,16,21). These fitness effects were linked to the absence of XA and increased levels of kynurenic acid (KYNA) (21). Due to the fitness cost associated with disrupting *kmo*, the researchers selected *cd* as an alternative target in subsequent gene drive studies (17). In that work *An. gambiae* homozygotes for a CRISPR/Cas9 gene drive inserted in *cd* did not display any apparent fitness costs (17,22,23), supporting its suitability as a target and model for population replacement gene drive systems.

In this study, we address the knowledge gap regarding the fitness effects of disrupting the *cd* gene in *An. stephensi* by analysing multiple knock-in and knock-out lines. We assessed key fitness parameters, including survival, fecundity, and midgut integrity. Our results show that while knock-out mutations in *cd* do not impose significant fitness costs, knock-in mutations involving the insertion of a fluorescent marker and sgRNA, result in reduced survival after blood feeding and decreased larva-to-adult survival in heterozygotes, indicating that the observed fitness effects are likely due to the transgene insertion rather than disruption of the *cd* gene itself.

## Results

### Knock-in but not knock-out mutations in *cd* impact survival and fertility

We investigated phenotypes observed in *cd* knock-out and knock-in transgenic lines which were generated for studying homing gene drives in *An. stephensi* (24). We assessed two knock-out lines (indicated with ^R^) which have indels at different positions in the *cd* gene (at the 225th and 384th amino acids of the protein), *cd*^*225R*^ and *cd*^*384R*^ and knock-in lines expressing a fluorescent marker and sgRNA (indicated with ^g^) inserted at the same positions, *cd*^*g225*^ and *cd*^*g384*^.

Larva-to-adult survival was assessed by rearing 200 larvae of each genotype for both the knock-in (*cd*^*g384*^) and knock-out (*cd*^*384R*^) lines (heterozygotes, homozygotes and their wildtype (WT) siblings) and monitoring the number of pupae that successfully eclosed into adults. Overall we found that the eclosion rate of heterozygous *cd*^*g384*^ females (50.3% [95%CI] = 0.45-0.55]) did not differ significantly from *cd*^*384R*^ females (OR = 1.22[0.62-1.08], Tukey HSD z = 1.42, p = 0.157) and wildtype females did not differ significantly from heterozygous females in either line (Tukey HSD all p > 0.05) (STable 1). Both male and female homozygotes of *cd*^*g384*^ failed almost entirely to eclose (>1% [0.2-2%]), while *cd*^*384R*^ did not differ significantly from heterozygotes (OR = 0.82, [95%CI] = [0.62-1.08], Tukey HSD z = 1.42, p = 0.33) or wildtype (OR = 0.78, [95%CI] = [0.59-1.03], Tukey HSD z = 1.77, p = 0.18).

To assess the fecundity and fertility of *cd*^*384R*^, *cd*^*g225*^ and *cd*^*g384*^, five replicate cages of ten females and ten males of each genotype were crossed to wildtype (SDA-500) mosquitoes in a 1:1 ratio. After blood feeding, females were separated into wells of EAgaL plates (25) where they could individually lay eggs, and we counted the total number of eggs laid per female, and what proportion of these hatched into L1 larvae. The *cd*^*g225*^ heterozygous females either did not feed during the experiment or died after taking a blood meal, making it impossible to generate *cd*^*g225*^ homozygotes, so only heterozygotes of this line could be assessed (STable 2). We also observed death after taking a blood meal in *cd*^*g384*^ homozygous females, but none of the other genotypes. This is investigated further in another experiment which specifically tracks survival after blood feeding.

We found that *cd*^*384R*^ female heterozygotes produced slightly more eggs (100, [95%CI] = 87.1-116) than *cd*^*g384*^ (94.4, [95%CI] = 80.1-111) (Tukey HSD; z = 0.571. p = 0.05), however there were no significant differences across lines or between male or female heterozygote crosses to SDA-500 mosquitoes. Crosses of *cd*^*384R*^ female homozygotes to SDA-500 males produced significantly more eggs on average (110 [91.1-132]) than the equivalent cross in *cd*^*g384*^ 66.5 [53.2-83.2] (z = 3.4, p <0.001). SDA-500 females mated to homozygous males from either *cd*^*384R*^ or *cd*^*g384*^ produced the lowest average egg counts (56.9 [43.4-74.7] and 66.7 [49.9-89.3] respectively) but this was not a statistically significant drop in fecundity for the equivalent crosses to heterozygous males (p > 0.05) (Fig. 1A, STable 3).

**Figure 1.**
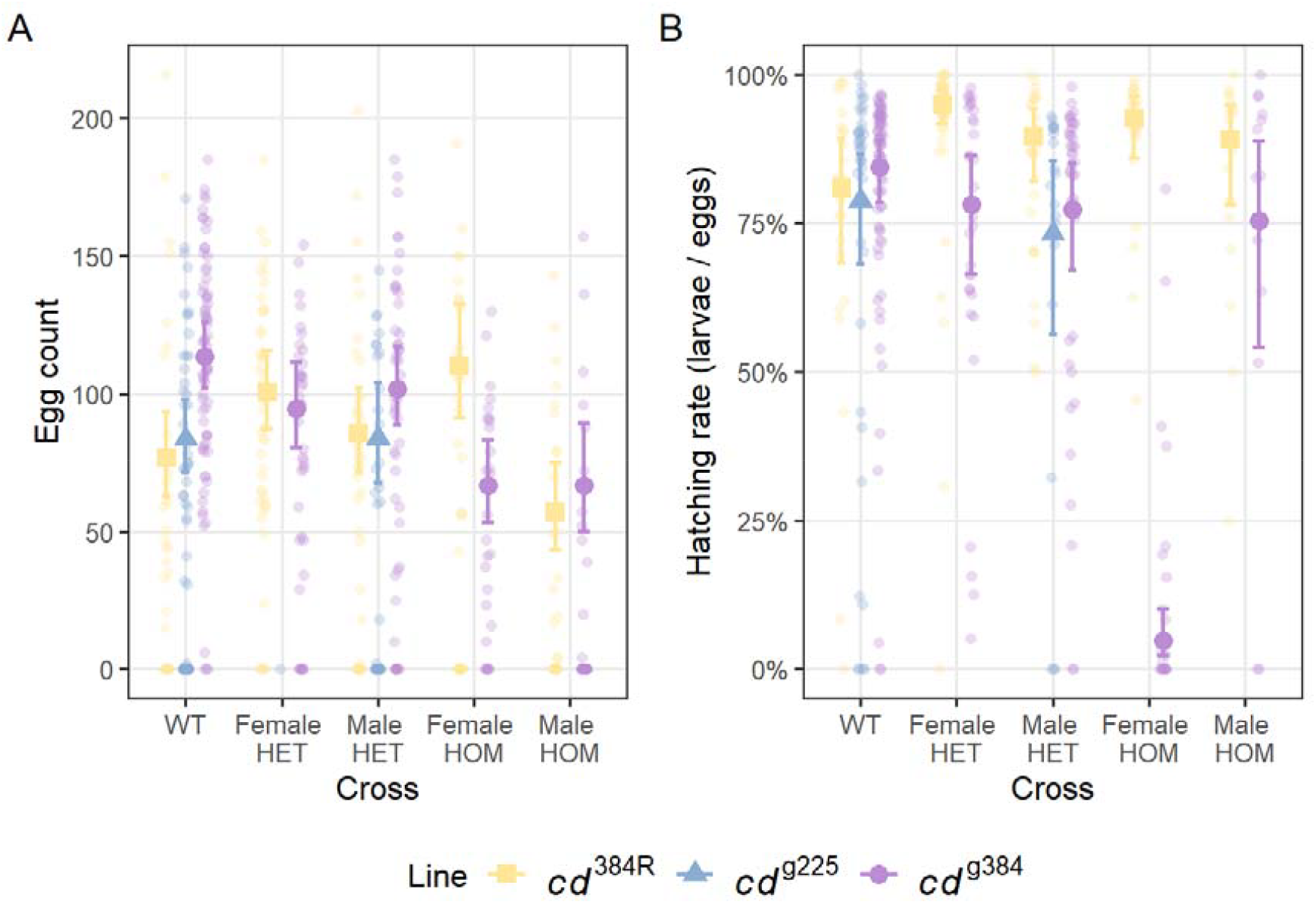
*cd*^*g384*^ homozygotes presented reduced reproductive fitness. Fecundity and fertility analysis of transgenic mosquitoes for the *cd*^*g225*^ (blue), *cd*^*g384*^ (purple), and *cd*^*384R*^ (yellow). (A) Fecundity was determined by the number of eggs laid per female. (B) Fertility was determined by the hatching rate of each female. WT, wildtype; HET, heterozygote; HOM, homozygote.

The *cd*^*384R*^ line had the highest overall hatching rates (proportion of eggs which successfully hatched) (heterozygote females = 0.95 [0.92 - 0.97]) and all crosses produced had either equivalent or slightly higher hatching rates than wildtype (Fig. 1B, STable 4). In contrast, we found an extremely poor hatching rate of *cd*^*g384*^ homozygous females crossed to SDA-500 males (0.05 [0.02 - 0.1]) (Tukey HSD p < 0.05 in all contrasts), indicating that *cd*^*g384*^ homozygous females are predominantly sterile. Other than this single directional cross, all other crosses of the *cd*^*g384*^ were not statistically significantly different from wildtype (Fig. 1B, STable 4). The only cross we were able to assess in the *cd*^*g225*^ line (male heterozygotes crossed to SDA-500 females) showed a marginally reduced (but not statistically significant) hatching rate compared to wildtype (0.73 [0.56 - 0.853], Tukey HSD z = 0.634, p = 0.52).

### Disruption of the *cd* gene by a knock-in results in death after a blood meal

To better assess the mortality after a blood meal observed in the fertility assay above, we again generated homozygous, heterozygous and wildtype females from the *cd*^*g384*^, *cd*^*g225*^ and *cd*^*225R*^ transgenic lines, as well as from the *cd*^*g384_del*^ and *cd*^*g338-384*^ knock-in lines previously generated for multiplexing studies in homing gene drives (24). The *cd*^*g384_del*^ line carries a single gRNA and fluorescent marker, with a deletion in the homology arms equivalent in size to the spacing of the four gRNAs expressed in *cd*^*g338-384*^ mosquitoes and so the disruption to the *cd* gene in these lines is equivalent. Mortality was monitored every 12h for five days following two separate blood meals. The survival after blood feeding of homozygous females of the *cd*^*g384_del*^, *cd*^*g338-384*^ and *cd*^*g225*^ knock-in lines was not assessed because heterozygous females from these lines died after a blood meal, hindering the generation of homozygotes.

All knock-in heterozygote lines showed substantially reduced survival relative to knock-out *cd*^*225R*^ homozygotes (85% survival [68.8 - 93.7] at 300 hours (h) post-blood feeding) (Fig. 2, STable 5). The *cd*^*g338-384*^ and *cd*^*g225*^ females had the most severe phenotypes, surviving only 7% and 9% as long as the knock-out *cd*^*225R*^ homozygotes (Time Ratio (TR) = 0.07, [95%CI] = [0.04-0.11], p < 0.001; TR = 0.09[0.06-0.14], p < 0.001), with 50% mortality at 58.9h [29.7-113.4] and 77.6h[40.1-150.7]) respectively. *cd*^*g384_del*^ heterozygous females also showed a 20% reduction in survival (TR = 0.20[0.12–0.32], p <0.001), with 50% mortality at 171h [87.2-335]. Although knock-in *cd*^*g384*^ heterozygous mosquitoes were similar to wildtype (TR = 1.49[0.93–2.36], p = 0.095), *cd*^*g384*^ homozygotes survived considerably less than knock-out *cd*^*225R*^ homozygotes (TR = 0.40[0.25–0.63], p <0.001). The wildtype siblings of all lines survived at least as long as the knock-out *cd*^*225R*^ homozygotes, if not better (STable 5). These results suggest that this phenotype is due to the transgene insertion instead of the disruption of the *cd* gene itself.

**Figure 2.**
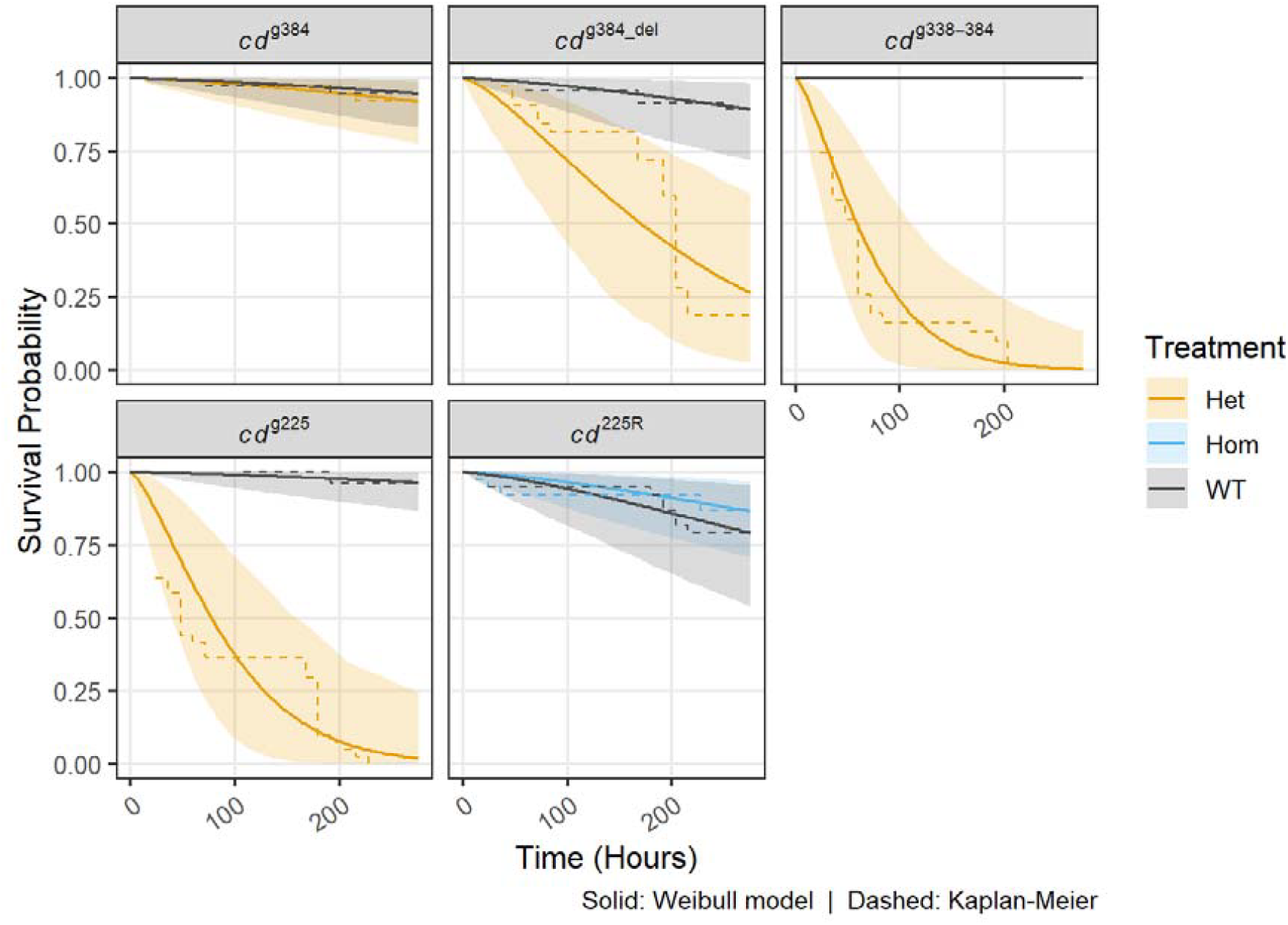
Knock-in lines expressing the Hr5IE1-ZsGreen fluorescent marker present a dominant fitness cost following a blood meal. Probability of survival after sequential blood-feeding of heterozygous (Het, orange) and/or homozygous (Hom, blue) females compared to their wildtype siblings (WT, gray). The survival after blood feeding is represented by the Kaplan-Meier survival curves (dashed line) and the Weibull parametric survival model (single solid line per genotype (n = 384)). Blood meals were provided at 0h and 168h. No measurements were taken between 120h and 168h.

### Mortality after a blood meal is associated with midgut barrier dysfunction

Previous work by others (21) has shown that mutants in the *kmo* gene, another gene in the ommochrome pathway, display similar mortality following a blood meal. This was linked to a disruption of the midgut barrier which could be determined by a ‘Smurf assay’. In this assay, a non-absorbable, non-toxic blue dye is administered through an artificial blood meal. Mosquitoes are scored as ‘Smurf positive’ if the dye leaks into the hemocoel and spreads throughout the body and cuticle within 24h, indicating midgut barrier disruption. In a ‘Smurf negative’ mosquito the blue dye stays in the digestive tract (SFig. 1).

In this experiment, we examined midgut barrier permeability in *cd*^*g225*^, *cd*^*225R*^, and wildtype females because the *cd*^*g225*^ knock-in line showed a severe phenotype. Wildtype mosquitoes had only a 5% probability of visible gut barrier disruption in our experiments (OR=0.06, [0.02 - 0.13], while *cd*^*g225*^ heterozygotes showed an elevated smurf rate of 79.8% [71-86.4] compared to wildtype controls (OR = 65.61[25.70–205.20], p <0.001), confirming that this knock-in insertion in the *cd* allele causes gut barrier disruption (Fig. 3B, STable 6). *cd*^*225R*^ homozygous mosquitoes (knock-out) showed no smurf positive individuals (Fig. 3B).

**Figure 3.**
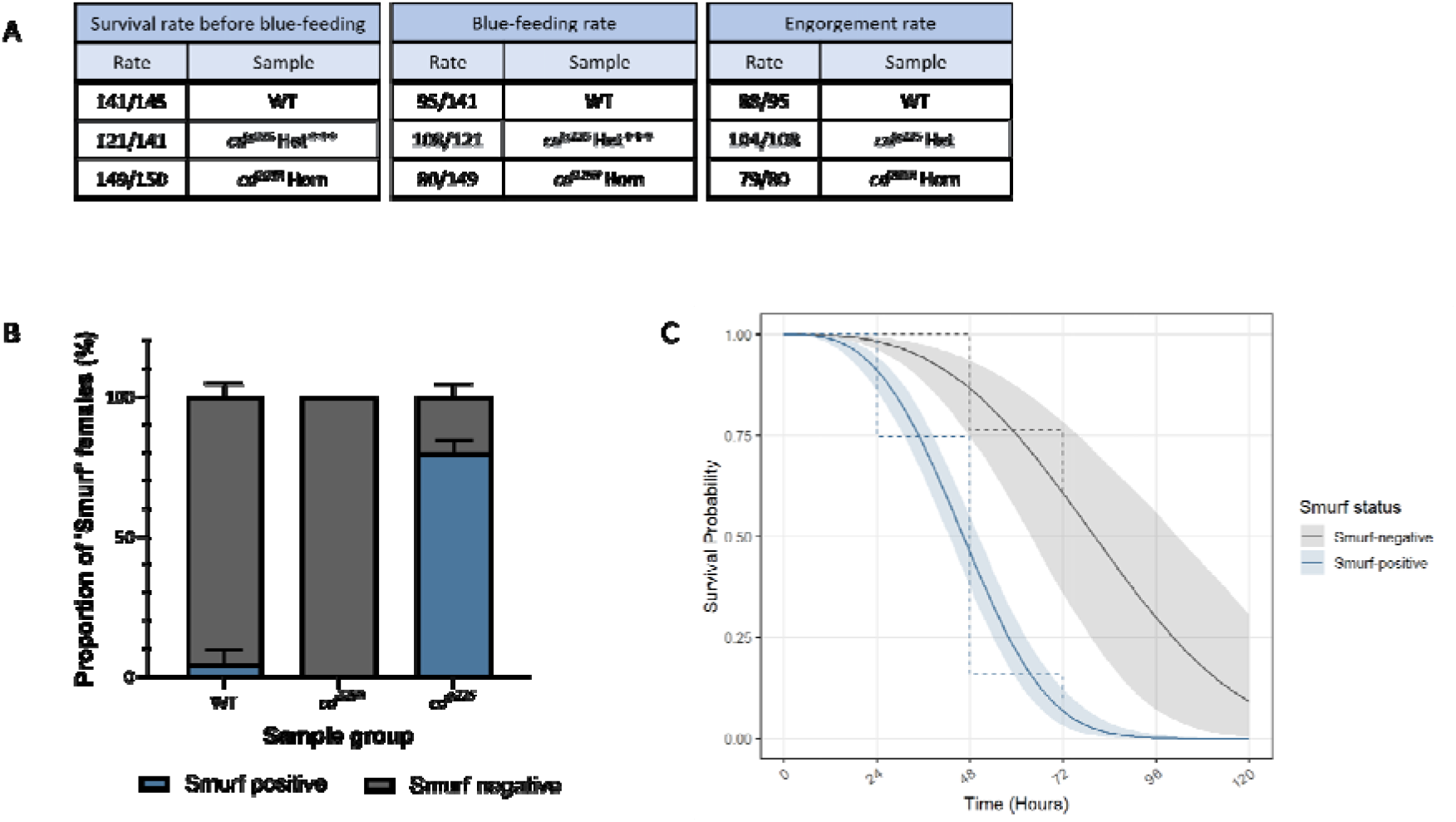
Mutations of the *cd gene by kno*ck-in result in a disruption of the integrity of the midgut barrier. (A) Strain performance at stages prior to ‘Smurf’ screening. A Fisher exact test of knock-in and knock-out in comparison to the control sample was performed at a 0.05 significance level; cells highlighted in light blue were significantly different from the control. (B) Proportion of ‘Smurf positive’ and ‘Smurf negative’ females one day after ‘blue-feeding’. The blue bar indicates the proportion of ‘Smurf positive’ and the grey bar the proportion of ‘Smurf negative’ females. Error bars represent the SEM. (C) Probability of survival after ‘blue-feeding’ of *cd*^*g225*^ heterozygotes according to their ‘Smurf’ status. The survival after blood feeding is represented by the Kaplan-Meier survival curves (dashed line) and the Weibull parametric survival model with a single solid line per genotype (n = 104). Het, heterozygote; Hom, homozygote; ***, p <0.001.

The survival after ‘blue-feeding’ was scored every 24h for a total of three days (Fig. 3C). All wildtype and homozygous knock-out *cd*^*225R*^ females survived up to 72h after the artificial blood meal (SFig. 2). However, for *cd*^*g225*^ knock-in heterozygous females, 72h survival was estimated at only 16.9% [0-0.8]. The estimate for wildtype and knock-out females was highly unstable due to complete or near-complete separation - no events (deaths) were observed in this group, rendering the time ratio unreliable (TR = 4900.99[0-∞], p = 0.998)). In conclusion, disruption of the *cd* gene through a transgene insertion but not by a knock-out mutation causes disruption of the midgut barrier, suggesting that the observed phenotype is more likely to be due to the transgene and not by disruption of the Cd protein (SFig. 3).

To determine whether the survival of *cd*^*g225*^ heterozygous females after ‘blue-feeding’ was correlated with their ‘Smurf’ score, the probability of survival was modelled using a Weibull parametric survival model using the ‘Smurf’ status as the sole factor (n = 104). At the 72h window, we found that Smurf positive females had a 0.06 [0.02-.11] probability of survival, while Smurf negative females had a greater than 50% chance of still being alive (0.61 [0.36-.78]). We estimated that ‘Smurf’ positive females had roughly half the survival time of ‘Smurf’ negative females (TR = 0.58[0.45-0.74], p <0.001) (Fig. 3C). This suggests that reduced probability of survival after blood feeding is partly driven by midgut barrier dysfunction; however, knock-in mosquitoes that do not show midgut barrier impairment (smurf negative) do still display mortality after a blood meal, indicating that additional factors are potentially involved or that the ‘Smurf’ assay is not sufficiently sensitive.

Additionally, knock-in females were significantly more likely to feed on the ‘Smurf’ solution compared to knock-out and wildtype females (Fig. 3A), and they remained fully engorged 24h after ‘blue-feeding’, whereas knock-out and wildtype females were able to digest and excrete the blue dye (SFig 1). These observations suggest that *cd*^*g225*^ heterozygous females may also have impaired sensory, digestive or excretory function, which could contribute to the increased mortality observed in this line regardless of their ‘Smurf’ phenotype.

### XA supplementation partially rescues the mortality after blood feeding phenotype

Regarding the mechanism underlying reduced survival after blood feeding phenotype, we hypothesised that disruption of the *cd* gene in *An. stephensi* may lead to the accumulation of 3-HK, a metabolite known to impair peritrophic matrix integrity in mosquitoes (3,26). We therefore tested whether oral supplementation with XA, an antioxidant, could mitigate 3-HK–associated damage and rescue the observed lethality phenotype in knock-in *cd*^*g225*^ heterozygotes as previously described in *kmo* knock-out mosquitoes (21).

We supplemented either through the sugar feeders, so that the adults were supplied with a constant supply of XA, or through a single dose with the blood meal (Fig. 4). The XA in blood treated group had significantly higher survival (83% [70.1-91%]) compared to the untreated (58% [41-73%]) at 72h post blood feeding; TR = 1.64[1.15–2.35], p = 0.007). The NaOH control group (6mM C−) did not significantly alter survival relative to the untreated group (TR = 0.9[0.68–1.19], p = 0.468), indicating that this effect is not attributable to the NaOH vehicle used to make the XA stock solution (Fig. 4, STable 7). Provision of XA via sugar feeding also increased survival of heterozygotes (94% [87.8-97.2%]) compared to unsupplemented (83.4% [72.7-90.6%]), although the overall survival in that experiment was also higher. Together, these results indicate that XA supplementation at 6mM significantly extends post-blood feeding survival in knock-in heterozygotes.

**Figure 4.**
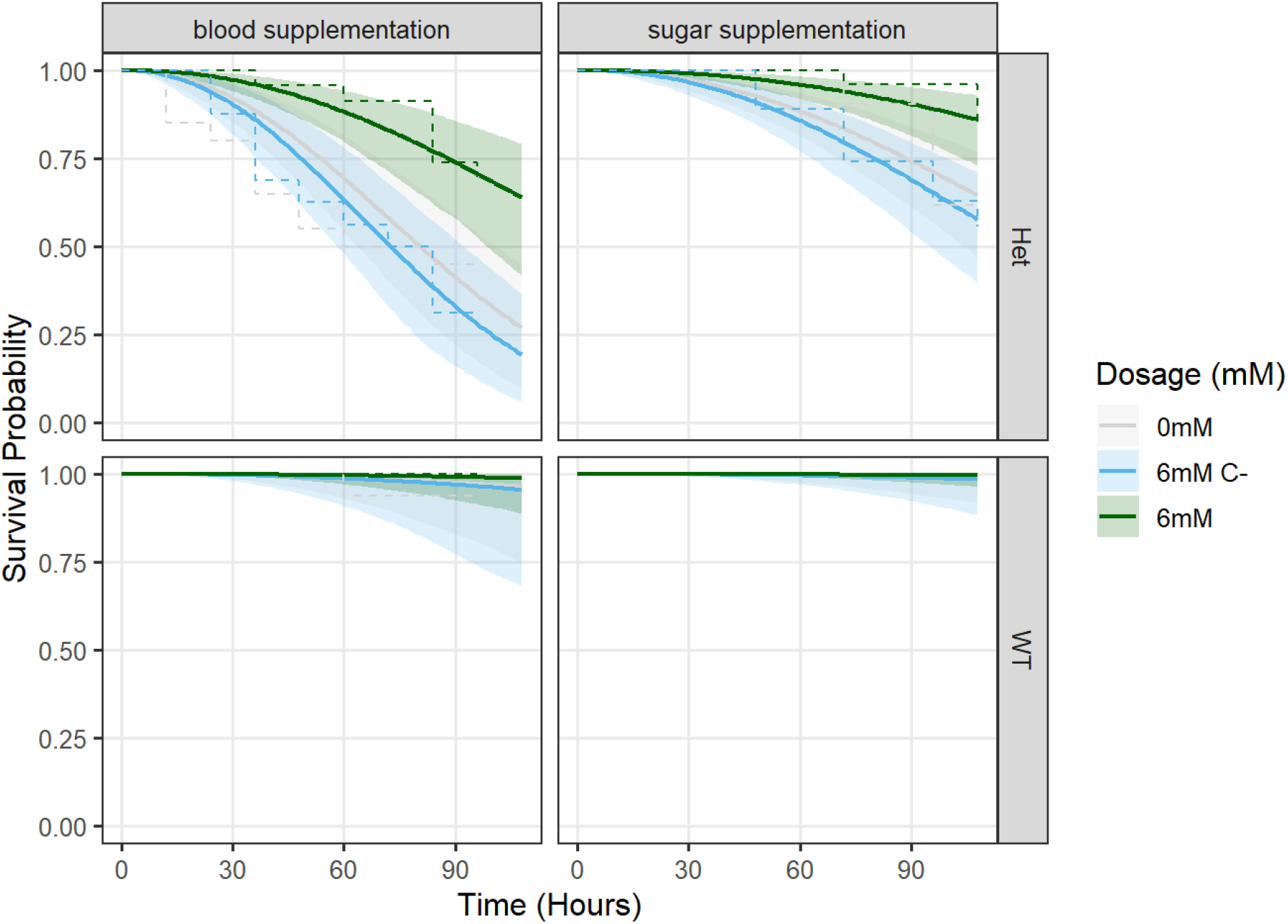
XA supplementation rescues the mortality after blood feeding phenotype in *cd*^*g225*^ heterozygotes. Probability of survival after blood feeding after oral administration of 6mM of XA (6mM, green) in comparison to its respective 0.01N NaOH control (C-, blue) and an untreated control (0mM, gray). Treatment was administered either through blood (left panels) or sucrose feeders (right panels). The number of dead females was counted every 12h after a blood meal. The data is presented by the Kaplan-Meier survival curves (dashed line) and a Weibull parametric survival model (solid line, n=387).

### *cd* expression is altered in the presence of the transgene

These results indicate that the phenotype observed in knock-in lines is linked to the presence of the inserted transgene rather than to disruption of the *cd* gene itself, as homozygous knock-outs do not exhibit reduced larva-to-adult survival, mortality after blood feeding, or midgut barrier dysfunction. To further assess whether the post-blood feeding survival phenotype is caused by disruption of *cd* function, we assessed the survival after blood feeding of *cd*^*g225*^*;cd*^*225R*^ trans-heterozygous females and compared it to *cd*^*g225*^ heterozygotes. Controls (*cd*^*225R*^ heterozygous and *cd*^*g225*^ sibling wildtype females) showed >98% survival across the experiment relative to the *cd*^*g225*^ heterozygotes (TR = 12218566.26, [95%CI] = [0.00–∞], p = 0.997 and TR = 26.66[11.04–64.38], p <0.001, STable 8) consistent with previous experiments (Fig. 5). However, the estimate of *cd*^*225R*^ heterozygotes is highly unstable, suggesting complete or near-complete separation in the data. *cd*^*g225*^ heterozygotes had an average survival probability of 20% [<0.001 - 0.61] at 100 hours, and *cd*^*225R*^ heterozygous mosquitoes did not differ significantly from this (TR = 0.97 [0.40–2.34], p = 0.944), indicating that the reported mortality after blood feeding phenotype is predominantly or solely linked to the insertion or the genetic background of the insertion (Fig. 5).

**Figure 5.**
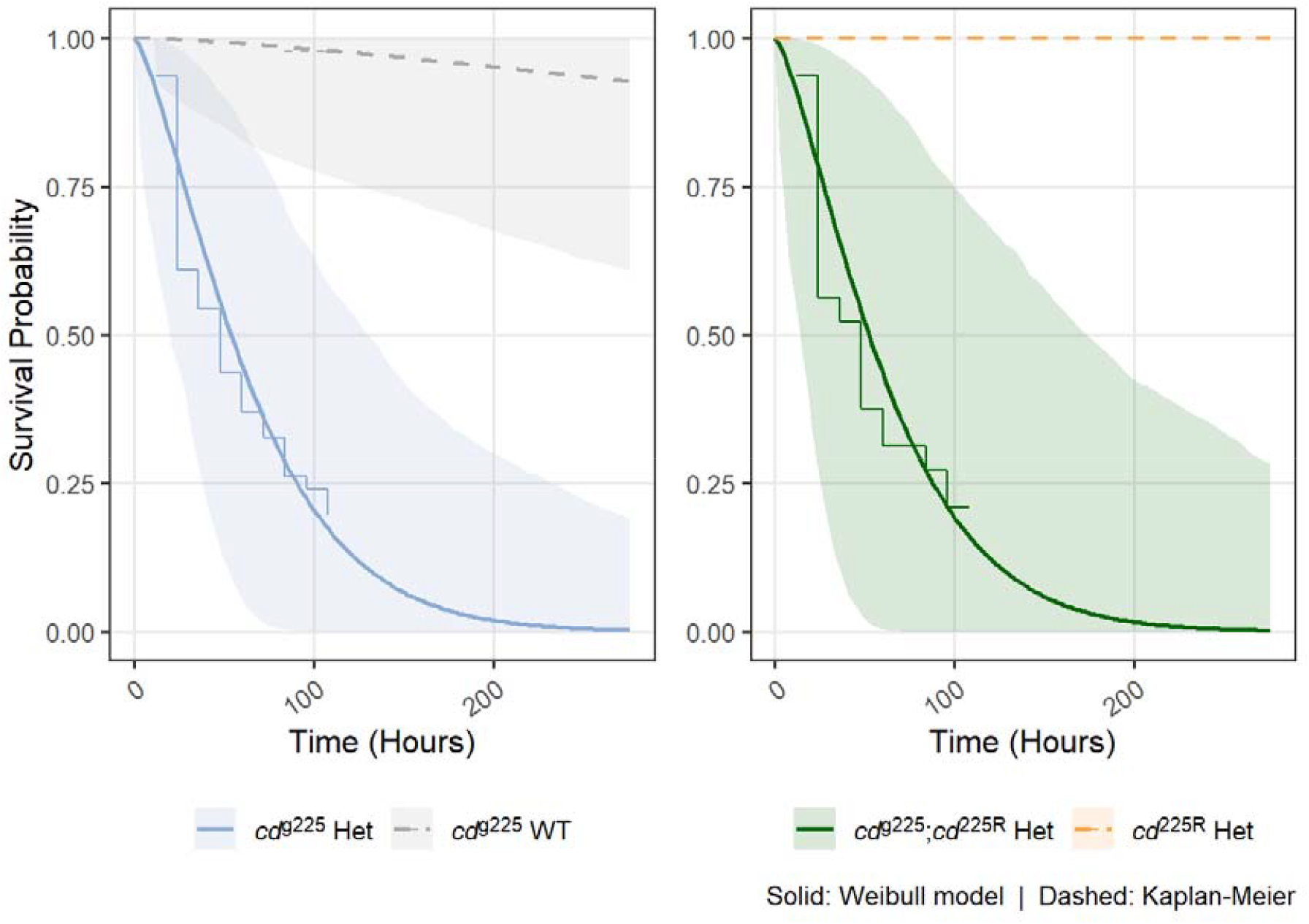
The probability of survival of *cd*^*g225*^;*cd*^*225R*^ trans-heterozygotes did not differ from *cd*^*g225*^ heterozygotes. Probability of survival after blood feeding of *cd*^*225R*^;*cd*^*g225*^ trans-heterozygous females in comparison to heterozygous females for both alleles and wildtype (WT) females. The survival after blood feeding is represented by the Kaplan-Meier survival curves (staircase-like) per genotype and by the Weibull parametric survival model with a curved continuous line (n = 384). The continuous lines represent genotypes that expressed the *cd*^*g225*^ transgene and the dashed lines indicate the absence of this allele. The time of the blood meal corresponds to 0h. The number of dead females was counted every 12h for a total of 5 days.

We hypothesised that the Hr5/IE1 promoter/enhancer element, which expresses the fluorescent protein marker in the transgenic construct, might alter splicing of the *cd* gene, as observed with similar knock-in lines (27). To investigate this, we performed 3’ RACE starting from the exon (Ex) upstream of the transgene insertion (Ex3), to characterise the transcripts produced. Sequencing of the 3’ RACE PCR amplicons revealed that both knock-in lines spliced correctly from Ex3 to Ex4, with transcription continuing into the transgene and terminating within the Hr5/IE1 promoter (SFig. 3). Analysis of the predicted amino acid sequence for the *cd*^*225R*^-derived transcript showed that following the sgRNA 225 cut site and a 1bp deletion the sequence continued for an additional 75 amino acids before reaching a stop codon (SFig. 3). Both the knock-in *cd*^*g225*^ and the knock-out *cd*^*225R*^ lines are predicted to generate similar truncated Cd proteins, suggesting that the pronounced fitness effects observed in the knock-in lines are primarily due to the transgene rather than disruption of the Cd protein. In contrast, the *cd*^*384R*^ line produces a nearly full-length Cd protein (with an in-frame deletion causing amino acids Q383 and A384 to be replaced by a single P residue), while the *cd*^*g384*^ line generates a truncated protein (SFig. 3), complicating direct fitness comparisons. Overall, these results indicate that canonical *cd* splicing is not disrupted, ruling out splicing defects as the cause of the observed phenotype. An alternative hypothesis to explain why the transgene could have an effect on fitness was that Hr5/IE1 was enhancing the expression of the truncated and putatively antimorphic or neomorphic *cd* allele and/or a nearby gene. To determine whether there were altered expression levels of the *cd* gene associated with the enhancing properties of the Hr5/IE1 promoter, a real-time qPCR was performed on the same samples used for the RACE analysis. We found *cd* expression levels were significantly downregulated in females and males of both knock-out lines (*cd*^*384R*^ and *cd*^*225R*^) (Fig. 6, SFig.4), except for *cd*^*225R*^ females when *cd* expression was normalised against *GAPDH* where it was not significant (permutation t-test, t = 0.797, df = 20, p = 0.077) (SFig. 4). In contrast, *cd* expression in females and males of the *cd*^*g225*^ was significantly higher than wildtype whereas *cd*^*g384*^ only showed a significant increase in relative expression in females when normalised against *rps7* (p <0.001) and in males when normalised against *GADPH* (p <0.001) (Fig. 6, SFig. 4). These results suggest that the different phenotype observed in the knock-in lines when compared to the knock-out lines might be attributed to the enhancing properties of Hr5/IE1. Moreover, the expression levels of *cd* were higher in females of the *cd*^*g225*^ line than in *cd*^*g384*,^ indicating that higher expression of Cd protein, or of the encoded mutant Cd protein, might be correlated with a lower probability of larva to adult survival and survival after a blood meal.

**Figure 6.**
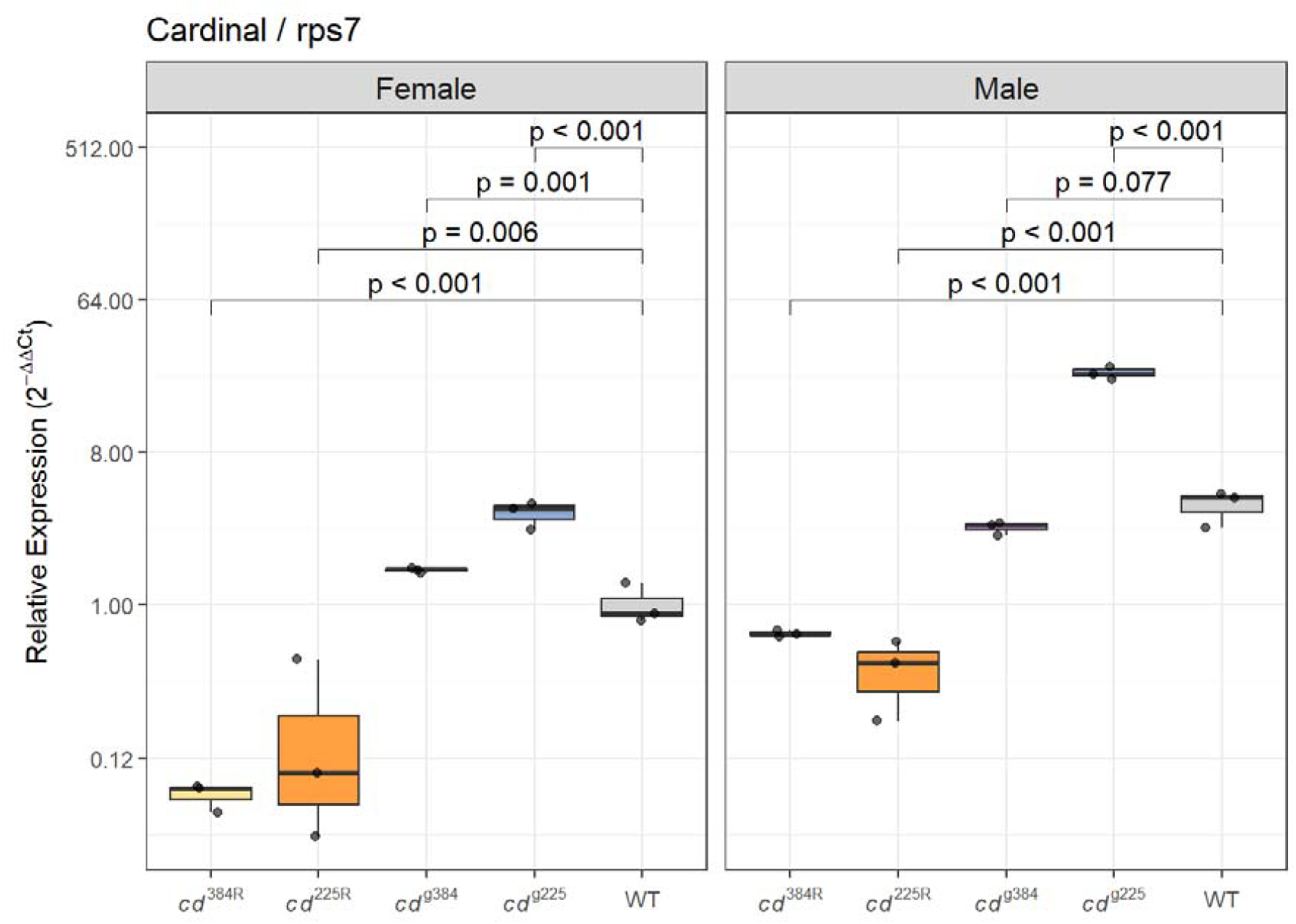
The Hr5/IE1 promoter enhances *cd* expression in the knock-in lines. Expression analysis of the cd gene by Taq-Man qPCR. Comparison of the expression levels of the cd gene of two different knock-in lines (*cd*^*g225*^ and *cd*^*g384*^) to their respective knock-out lines (*cd*^*225R*^ and *cd*^*384R*^) and using the wildtype (WT) female sample as a reference. cd expression was normalised against rps7 (housekeeping gene). The relative expression (RQ) is plotted on a log2-scaled axis. A permuted t-test with a 0.05 significance level was performed on raw deltaCT values for each line in comparison to the control sample to determine statistical significance.

## Discussion

Here we present evidence that the use of the Hr5/IE1 promoter in transgenic constructs in *An. stephensi* is associated with unintended fitness costs and upregulated gene expression. In previous work, we developed several knock-in and knock-out lines targeting the *cd* gene. We noticed that although homozygotes (or transheterozygotes) of both types of mutations displayed the expected eye-colour phenotype, knock-in mutant females, even as heterozygotes, would often die after taking a blood meal and had decreased fertility compared to the knock-out lines. Previous reports in *An. stephensi* observed similar fitness defects in mutants of another gene in the ommochrome pathway, *kmo*. In that report, death after blood feeding was observed in knock-out mutants of three mosquito species. This was found to be a result of metabolic imbalances, which resulted in midgut barrier dysfunction, with accumulation of kynurenic acid, the precursor to 3-HK, the root cause. Supplementation with XA (the downstream metabolite after 3-HK) rescues the phenotype. It was suggested in that work, and others, that *cd* (PHS) was not directly along this pathway, rather that it was part of a reaction converting 3-HK to ommochrome (18,21). It was thus expected that *cd* mutants would not share this same phenotype.

Initial generation of *cd* knock-out mutants agreed with this suggestion, with normal fertility and no mortality after taking blood meals, these lines are maintained as homozygotes without issue. However, after generating the knock-in lines, we observed fitness defects, initially with females from some lines dying after taking a blood meal. When generating homozygotes of these lines in order to measure their fitness, we found fertility defects. We then investigated these observations more systematically.

We found that our knock-in lines had unexpected fitness costs, more significantly when they were homozygous. *cd*^*g384*^ had the least fitness defects of the knock-in lines, though females of the *cd*^*g384*^ line had reduced fertility. Post-blood meal survival of heterozygous females of three other knock-in lines *cd*^*g384_del*^, *cd*^*g338-384*^ and *cd*^*g225*^ was so strongly reduced that it was difficult to reliably generate enough homozygotes for experimentation. We performed a ‘Smurf’ assay to examine midgut barrier integrity in one such line, *cd*^*g225*^ and found that the majority of the females showed disruption of the midgut and reduced survival after ‘blue-feeding’. Supplementation with XA either in the sugar or blood meals could partially rescue the females, increasing the longevity after a blood meal, although not to wildtype levels. Synthesis of XA is dramatically increased after a blood meal, reaching its peak at 24h after the ingestion of blood and controlling the oxidative stress in the midgut (3). XA is known for its role in maintaining the midgut epithelial homeostasis and as an endogenous antioxidant in both midgut and malpighian tubules through its ability to bind to both heme and/or iron, which mainly occurs at a slightly alkaline pH (3). It may be that the rescue effect observed was at least partially due to the antioxidant properties of XA, rather than indicating a role for *cd* in this part of the pathway. In *kmo* knock-out females, low survival after a blood meal was partially reverted by XA supplementation, in that case not through its oxidative role but due to bacterial proliferation in the gut (21). Further experiments are necessary to further investigate the role of *cd* in this pathway in *An. stephensi*.

A cross between knock-in and knock-out mosquitoes revealed that the fitness effect is associated with carrying the transgene. 3’RACE confirmed that splicing was not disrupted in the knock-in lines and that both knock-in and knock-out lines produce similar truncated proteins. qPCR showed that *cd* transcripts were overexpressed in the knock-in lines, which we hypothesize is due to the AcMNPV Hr5/IE1 promoter/enhancer expressing the fluorescent protein marker. Although this Baculovirus-derived promoter has been widely used in various insect species without reported deleterious effects (28–30), a recent study in *An. gambiae* found that constructs using a variant of another Baculovirus promoter, OpIE2a, for fluorescent protein expression caused similar phenotypes (31). In that study, females failed to blood feed due to a morphological defect in the proboscis, which impaired feeding. These findings underscore the importance of carefully evaluating the effects of commonly used transgenic construct components, especially when applied in new species or construct contexts.

Following the results found here further studies to characterise the expression levels of the *HKT* gene and the concentration levels of XA and 3-HK at different mosquito stages would be key to better understand the role of this protein in the regulation of the synthesis of XA, if any. Moreover, better understanding of the regulation of the timing of XA synthesis could be a potential tool for vector control. Targeting a gene involved in the regulation of XA synthesis might yield a late acting female-specific phenotype as a result of oxidative stress which would be of interest for the control of vector populations.

## Materials and Methods

### Mosquito strains and lines

The *An. stephensi* SDA-500 strain was used as the wildtype strain. The mutant lines used in this study (*cd*^*g225*^, *cd*^*g384*^, *cd*^*g384_del*^, *cd*^*g338-384*^, *cd*^*225R*^, and *cd*^*384R*^) were generated in previous work using CRISPR/Cas9 (24,32). All experiments performed for this study were approved by the Biological Agents and Genetic Modification Safety Committee of The Pirbright Institute and/or the Biological Agents Safety Committee of The University of York Biology Department and undertaken in accordance with all relevant ethical regulations.

### Maintenance of mosquito colony

*Anopheles stephensi* SDA-500 mosquitoes were maintained under standard conditions at 70-80% relative humidity, 28±1 °C, and 14:10h day-night cycle (27,33). Larvae were reared in pools of 200 and fed with Sera Micron (Olibetta) at the first instar stage and with Extra Select Pond Pellets Complete Fish Food at later stages. Adults were maintained in cages where they were provided 10% sucrose *ad libitum* and females were offered a blood meal of defibrinated horse blood (TCS Biosciences) delivered with a Hemotek membrane feeding system (Hemotek, Inc) covered with a double layer of Parafilm (Bemis).

### Phenotype analysis

To determine the larva to adult survival, homozygous, heterozygous and wildtype mosquitoes from the *cd*^*g384*^ and *cd*^*384R*^ transgenic lines were reared under standard rearing conditions and screened as late larvae using a fluorescent stereo microscope (Leica M165FC) to determine the genotype ratios. Heterozygous larvae were identified by the presence of fluorescence, and homozygous larvae exhibited both fluorescence and a pink-eyed phenotype. Wildtype larvae lack both traits. The number of females and males reaching pupal and adult stages were recorded to calculate the probability of survival of each genotype.

The fecundity and fertility of wildtype, heterozygotes and/or homozygotes from the *cd*^*g225*^, *cd*^*g384*^, and *cd*^*384R*^ transgenic lines were assessed using EAgaL plates (25) following the protocol described in (27).

For the survival after blood feeding assay, pools of 10 wildtype, heterozygous and/or homozygous females were separately crossed to males of the SDA-500 strain in a 1:1 ratio for a total of five replicates per genotype. Females were allowed to mate for three days when they were offered a blood meal for 90min. 8h post-blood meal, non-engorged females were counted and removed from each replicate. The number of dead females was scored every 12h for a total of 5 days. A week after the first blood meal, the remaining females were blood fed for a second time and the consequent survival was scored. Non-engorged females were also removed after the second blood meal.

### Smurf assay

L1 larvae from *cd*^*g225*^, *cd*^*225R*^ and wildtype maintenance cages were sorted using a Biosorter (Union Biometrica) and reared under standard conditions. 50 females from each genotype were screened, sexed and crossed as adults to wildtype males in a 1:1 ratio for a total of three replicates. 7 days post-mating, females were offered an artificial feeding solution prepared using 150mM of NaCl (5M NaCl, ThermoFisher Scientific - #AM9760G) and 10mM of NaHCO_3_ (5% NaHCO_3_ aqueous solution, Electron Microscope Sciences - #26105-02), and supplemented with 2.5% of a 320g/L stock solution of FD&C blue dye (Erioglaucine disodum salt, Sigma-Aldrich - #861146) as previously described by (21). The resulting solution was named ‘blue-feeding’ solution. Non-engorged females were counted and removed from the cages. 20-28h post feeding, females were screened for the Smurf phenotype using a Leica M165FC microscope. A female was scored as ‘Smurf negative’ when blue coloration was detected only within the digestive tract whereas ‘Smurf positive’ females presented blue coloration spread throughout the abdominal cavity, cuticle, and legs (SFig. 1). ‘Smurf positive’ and ‘Smurf negative’ females per genotype were reared in separate cages to assess their survival every 24h for a duration of 3 days.

### Expression and splicing of the *cd* gene

The RNA of pooled *cd*^*g384*^ and *cd*^*g225*^ heterozygous, *cd*^*384R*^ and *cd*^*225R*^ homozygous, and wildtype females was extracted using the NucleoSpin RNA kit (Macherey-Nagel). The SMARTer RACE 5’/3’ Kit (Takara Bio) was used to generate the RACE ready cDNA and for the RACE PCR with gene specific primers: LA8430: 5’CCGCCTCTCGATGGGGCCAATGC and nested primer LA8431: 5’CGACGGCGGCGGATTTCCCTGAG. For qRT-PCR the LunaScript RT SuperMix Kit (New England Biolabs) was used to synthesise the cDNA and assembled in the qPCR reaction with the Luna Universal Probe qPCR Master Mix (New England Biolabs). TaqMan primer/probe mixes were designed and synthesized by ThermoFisher Scientific; *cd* (ASTE006748) Assay ID: APXGZTR; *rsp*7 (ASTE004816) Assay ID: APKCDR; *GAPDH* (ASTE006277) Assay ID: APH6H6W.

### Statistical analysis

All analyses were performed in R (v4.5.2)(34). Fecundity (egg counts) was modelled using a zero-inflated negative binomial mixed model fitted with glmmTMB, with cross direction and genetic line as fixed effects and replicate and plate well as nested random effects. Hatching rate was modelled using a binomial generalised linear mixed model (GLMM), also fitted with glmmTMB (35). Estimated marginal means and pairwise contrasts with family-wise error rates were obtained using emmeans. Post-blood feeding survival, trans-heterozygote survival, and XA supplementation survival data were modelled using Kaplan Meier estimates alongside Weibull parametric survival models fitted with flexsurv (36). Midgut integrity (smurf assay) and adult eclosion data were modelled using binomial generalised linear models (GLMs). Gene expression data were analysed using the delta-delta-Ct method; ΔCt levels were modelled with a permutation-based linear model using emmeans for contrasts (37). Model tables were produced using sjPlot. All figures were produced using ggplot2 (38).

## Data availability

Raw data and supporting code are available at https://github.com/Philip-Leftwich/cardinal_fitness At submission this will be deposited into an open access repository to produce a permanent doi.

## Author contributions

MLG, LS, EG, JA, ME, KN, JL, MM performed the crosses and collected inheritance data. PTL and MLG performed statistical analysis. MA, RN and LA supervised the work. MLG and LS wrote the initial draft and MAEA, PTL, EG and LA revised the manuscript. LA procured funding and MLG, EG, MA, and LA conceptualised the work. All authors read and approved the manuscript.

## References

1. Han Q, Beerntsen BT, Li J. The tryptophan oxidation pathway in mosquitoes with emphasis on xanthurenic acid biosynthesis. J Insect Physiol. 2007 Mar 1;Physiology of Vector Arthropods53(3):254–63. doi:10.1016/j.jinsphys.2006.09.004

2. Hebbar S, Traikov S, Hälsig C, Knust E. Modulating the Kynurenine pathway or sequestering toxic 3-hydroxykynurenine protects the retina from light-induced damage in Drosophila. PLOS Genet. 2023 Mar 23;19(3):e1010644. doi:10.1371/journal.pgen.1010644

3. Lima VLA, Dias F, Nunes RD, Pereira LO, Santos TSR, Chiarini LB, et al. The Antioxidant Role of Xanthurenic Acid in the Aedes aegypti Midgut during Digestion of a Blood Meal. PLOS ONE. 2012 Jun 11;7(6):e38349. doi:10.1371/journal.pone.0038349

4. Cerstiaens A, Huybrechts J, Kotanen S, Lebeau I, Meylaers K, De Loof A, et al. Neurotoxic and neurobehavioral effects of kynurenines in adult insects. Biochem Biophys Res Commun. 2003 Dec 26;312(4):1171–7. doi:10.1016/j.bbrc.2003.11.051

5. Oxenkrug GF, Navrotskaya V, Voroboyva L, Summergrad P. Extension of life span of Drosophila melanogaster by the inhibitors of tryptophan-kynurenine metabolism. Fly (Austin). 2011 Oct 1;5(4):307–9. doi:10.4161/fly.5.4.18414

6. Savitz J. The kynurenine pathway: a finger in every pie. Mol Psychiatry. 2020 Jan;25(1):131–47. doi:10.1038/s41380-019-0414-4

7. Lugo-Huitrón R, Ugalde Muñiz P, Pineda B, Pedraza-Chaverrí J, Ríos C, Pérez-de la Cruz V. Quinolinic Acid: An Endogenous Neurotoxin with Multiple Targets. Oxid Med Cell Longev. 2013;2013(1):104024. doi:10.1155/2013/104024

8. Garcia GE, Wirtz RA, Barr JR, Woolfitt A, Rosenberg R. Xanthurenic Acid Induces Gametogenesis in Plasmodium, the Malaria Parasite*. J Biol Chem. 1998 May 15;273(20):12003–5. doi:10.1074/jbc.273.20.12003

9. Maciel LG, Ferraz MVF, Oliveira AA, Lins RD, dos Anjos JV, Guido RVC, et al. Inhibition of 3-Hydroxykynurenine Transaminase from Aedes aegypti and Anopheles gambiae: A Mosquito-Specific Target to Combat the Transmission of Arboviruses. ACS Bio Med Chem Au. 2023 Apr 19;3(2):211–22. doi:10.1021/acsbiomedchemau.2c00080

10. Beard CB, Benedict MQ, Primus JP, Finnerty V, Collins FH. Eye Pigments in Wild-Type and Eye-Color Mutant Strains of the African Malaria Vector Anopheles gambiae [Internet]. [cited 2026 Apr 21]. Available from: 10.1093/oxfordjournals.jhered.a111606

11. Li J, Beerntsen BT, James AA. Oxidation of 3-hydroxykynurenine to produce xanthommatin for eye pigmentation: a major branch pathway of tryptophan catabolism during pupal development in the Yellow Fever Mosquito, Aedes aegypti. Insect Biochem Mol Biol. 1999 Apr 1;29(4):329–38. doi:10.1016/S0965-1748(99)00007-7

12. Wiley K, Forrest HS. Terminal synthesis of xanthommatin in Drosophila melanogaster. IV. Enzymatic and nonenzymatic catalysis. Biochem Genet. 1981 Dec;19(11–12):1211–21. doi:10.1007/BF00484574 PubMed PMID: 6802132.

13. Hebbar S, Traikov S, Hälsig C, Knust E. Modulating the Kynurenine pathway or sequestering toxic 3-hydroxykynurenine protects the retina from light-induced damage in Drosophila. PLOS Genet. 2023 Mar 23;19(3):e1010644. doi:10.1371/journal.pgen.1010644

14. Li M, Akbari OS, White BJ. Highly Efficient Site-Specific Mutagenesis in Malaria Mosquitoes Using CRISPR. G3 GenesGenomesGenetics. 2018 Feb 1;8(2):653–8. doi:10.1534/g3.117.1134

15. Heu CC, Gross RJ, L. KP, LeRoy DM, Fan B, Hull JJ, et al. CRISPR-mediated knockout of cardinal and cinnabar eye pigmentation genes in the western tarnished plant bug. Sci Rep. 2022 Mar 22;12(1):4917. doi:10.1038/s41598-022-08908-4

16. Gantz VM, Jasinskiene N, Tatarenkova O, Fazekas A, Macias VM, Bier E, et al. Highly efficient Cas9-mediated gene drive for population modification of the malaria vector mosquito Anopheles stephensi. Proc Natl Acad Sci. 2015 Dec 8;112(49):E6736–43. doi:10.1073/pnas.1521077112

17. Carballar-Lejarazú R, Ogaugwu C, Tushar T, Kelsey A, Pham TB, Murphy J, et al. Next-generation gene drive for population modification of the malaria vector mosquito, Anopheles gambiae. Proc Natl Acad Sci. 2020 Sep 15;117(37):22805–14. doi:10.1073/pnas.2010214117

18. Xu X, Harvey-Samuel T, Yang J, Alphey L, You M. Ommochrome pathway genes kynurenine 3-hydroxylase and cardinal participate in eye pigmentation in Plutella xylostella. BMC Mol Cell Biol. 2020 Sep 11;21(1):63. doi:10.1186/s12860-020-00308-8

19. Alphey L. Genetic Control of Mosquitoes. Annu Rev Entomol. 2014 Jan 7;59(1):205–24. doi:10.1146/annurev-ento-011613-162002

20. Raban R, James AA, Akbari OS. Advances in CRISPR gene drives for mosquito population control. Curr Opin Microbiol. 2026 Apr 1;90:102712. doi:10.1016/j.mib.2026.102712

21. Bottino-Rojas V, Ferreira-Almeida I, Nunes RD, Feng X, Pham TB, Kelsey A, et al. Beyond the eye: Kynurenine pathway impairment causes midgut homeostasis dysfunction and survival and reproductive costs in blood-feeding mosquitoes. Insect Biochem Mol Biol. 2022 Mar 1;142:103720. doi:10.1016/j.ibmb.2022.103720

22. Carballar-Lejarazú R, Dong Y, Pham TB, Tushar T, Corder RM, Mondal A, et al. Dual effector population modification gene-drive strains of the African malaria mosquitoes, Anopheles gambiae and Anopheles coluzzii. Proc Natl Acad Sci. 2023 Jul 18;120(29):e2221118120. doi:10.1073/pnas.2221118120

23. Carballar-Lejarazú R, Tushar T, Pham TB, James AA. Cas9-mediated maternal effect and derived resistance alleles in a gene-drive strain of the African malaria vector mosquito, Anopheles gambiae. Genetics. 2022 Jun 1;221(2):iyac055. doi:10.1093/genetics/iyac055

24. Larrosa-Godall M, Shackleford L, Edgington MP, Leftwich PT, Luk JCY, Southworth J, et al. Integrating multiplexing into confineable gene drives effectively overrides resistance in Anopheles stephensi. Nat Commun. 2026 May 7. doi:10.1038/s41467-026-72835-5

25. Tsujimoto H, Adelman ZN. Improved Fecundity and Fertility Assay for Aedes aegypti using 24 Well Tissue Culture Plates (EAgaL Plates). J Vis Exp JoVE. 2021 May 4;(171):e61232. doi:10.3791/61232

26. Feng Y, Peng Y, Song X, Wen H, An Y, Tang H, et al. Anopheline mosquitoes are protected against parasite infection by tryptophan catabolism in gut microbiota. Nat Microbiol. 2022 May;7(5):707–15. doi:10.1038/s41564-022-01099-8

27. Larrosa-Godall M, Ang JXD, Leftwich PT, Gonzalez E, Shackleford L, Nevard K, et al. Challenges in developing a split drive targeting dsx for the genetic control of the invasive malaria vector Anopheles stephensi. Parasit Vectors. 2025 Feb 7;18(1):46. doi:10.1186/s13071-025-06688-0

28. Anderson MAE, Gonzalez E, Edgington MP, Ang JXD, Purusothaman DK, Shackleford L, et al. A multiplexed, confinable CRISPR/Cas9 gene drive can propagate in caged Aedes aegypti populations. Nat Commun. 2024 Jan 25;15(1):729. doi:10.1038/s41467-024-44956-2

29. Grossman GL, Rafferty CS, Clayton JR, Stevens TK, Mukabayire O, Benedict MQ. Germline transformation of the malaria vector, Anopheles gambiae, with the piggyBac transposable element. Insect Mol Biol. 2001;10(6):597–604. doi:10.1046/j.0962-1075.2001.00299.x

30. Meredith JM, Underhill A, McArthur CC, Eggleston P. Next-Generation Site-Directed Transgenesis in the Malaria Vector Mosquito Anopheles gambiae: Self-Docking Strains Expressing Germline-Specific phiC31 Integrase. PLOS ONE. 2013 Mar 13;8(3):e59264. doi:10.1371/journal.pone.0059264

31. Sarig A, Sar-Shalom E, Kolley ESM, Yonah ES, Lamdan LB, Lewin AR, et al. An OpIE2-DsRed marker disrupts female blood-feeding and shortens lifespan in the malaria vector Anopheles gambiae. Genetics. 2025 Dec 31;iyaf280. doi:10.1093/genetics/iyaf280

32. Gonzalez E, Anderson MAE, Ang JXD, Nevard K, Shackleford L, Larrosa-Godall M, et al. Optimization of sgRNA expression with RNA pol III regulatory elements in Anopheles stephensi. Sci Rep. 2025 Apr 18;15(1):13408. doi:10.1038/s41598-025-98557-0

33. Southworth J, Gonzalez E, Nevard K, Larrosa-Godall M, Alphey L, Anderson MAE. Expanding the transgene expression toolbox of the malaria vector Anopheles stephensi. Insect Mol Biol. 2025;34(1):104–10. doi:10.1111/imb.12953

34. R Core Team. R: A language and Environment for Statistical Computing [Internet]. VIenna, Austria: R Foundation for Statistical Computing; 2023. Available from: https://www.R-project.org/

35. Brooks M E, Kristensen K, Benthem K J ,van Magnusson A, Berg C W, Nielsen A, et al. glmmTMB Balances Speed and Flexibility Among Packages for Zero-inflated Generalized Linear Mixed Modeling. R J. 2017;9(2):378. doi:10.32614/RJ-2017-066

36. Jackson C. flexsurv: A Platform for Parametric Survival Modeling in R. J Stat Softw. 2016 May 12;70:1–33. doi:10.18637/jss.v070.i08

37. Russell V. Lenth, Julia Piaskowski. Estimated Marginal Means, aka Least-Squares Means [R package version 2.0.3] [Internet]. 2026. Available from: https://rvlenth.github.io/emmeans/

38. Wickham H. ggplot2 [Internet]. Cham: Springer International Publishing; 2016 [cited 2024 Oct 14]. (Use R!). Available from: http://link.springer.com/10.1007/978-3-319-24277-4 doi:10.1007/978-3-319-24277-4

