## Supplementary information for "Knock-in ≠ knock-out: differential fitness effects of *cardinal* mutations in *Anopheles stephensi*"

**Supplemental information**

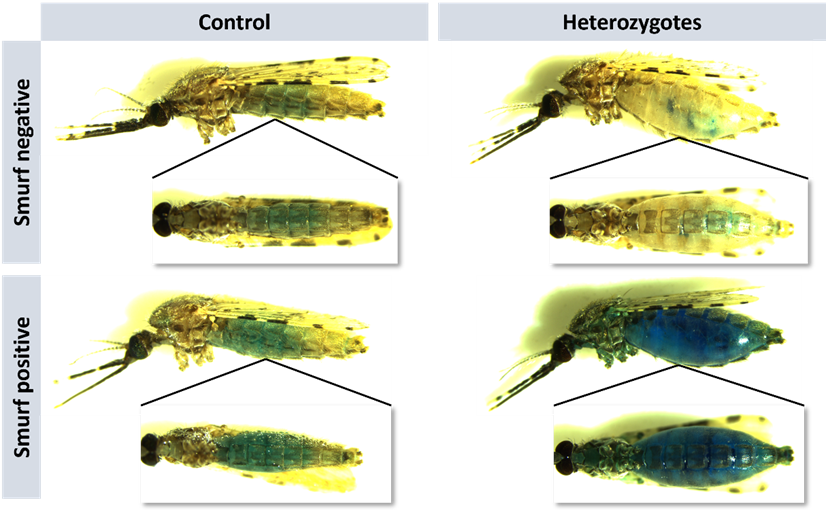

**SFig 1. Midgut permeability (‘Smurf assay’) in *cd^g225^* heterozygous and SDA-500 (control) adult females.** Representative images of heterozygous knock-in and WT adult female mosquitoes showing normal or loss of midgut integrity. Images were taken 20-22h post ‘blue-feeding’. For the lateral images, the exposure was set at 117.74ms, the gain at 1, and the magnification at 1.1x. The exposure was decreased to 150.51ms in ventral images and the magnification was increased to 1.4x. All the images are taken at a 70% of white light intensity in a Leica DFC7000T camera assembled on a Leica M165FC microscope.

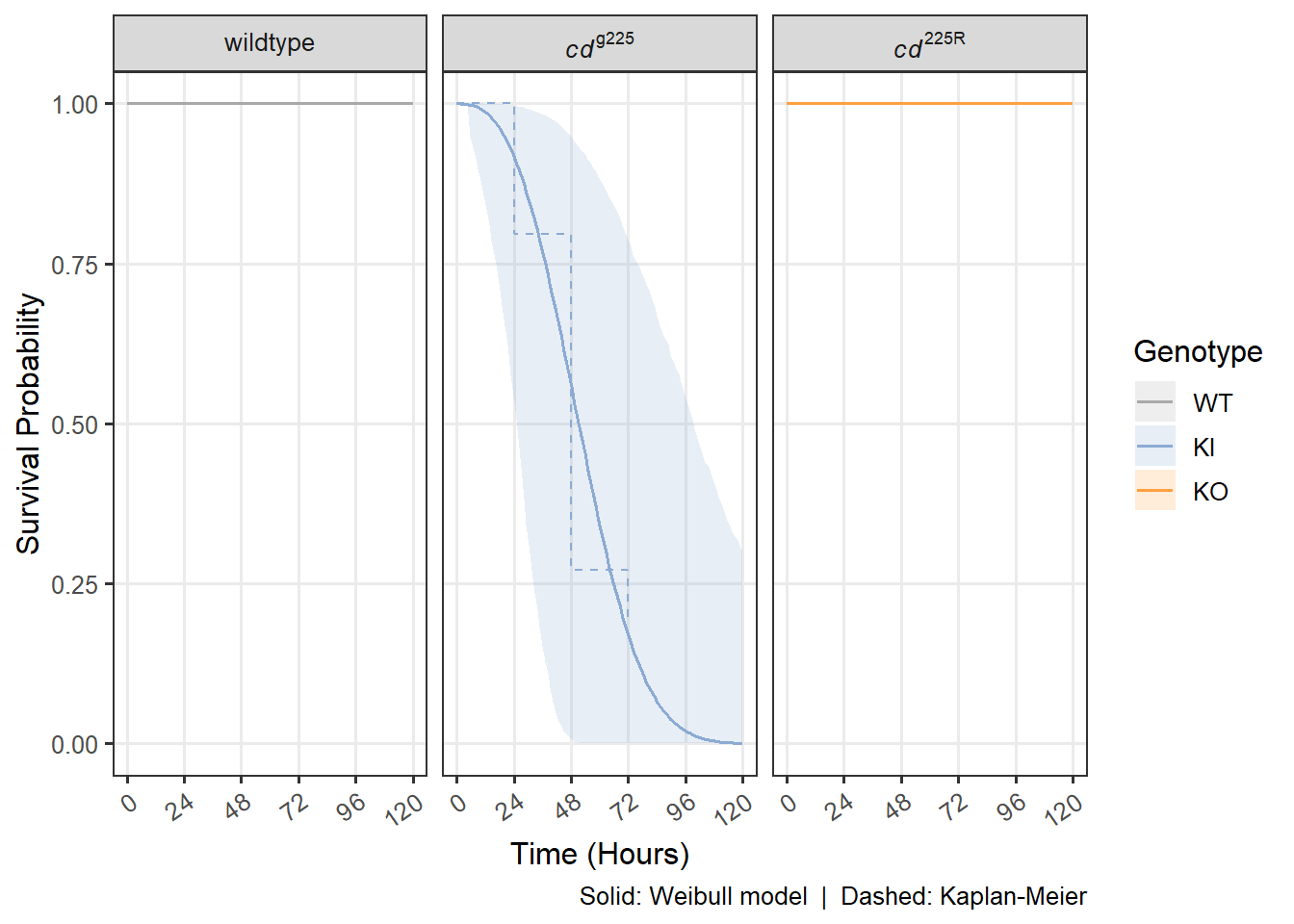

**SFig 2. Only *cd^g225^* heterozygous mosquitoes presented a low probability of survival after ‘blue-feeding’.** Probability of survival after ‘blue-feeding’ of WT (gray), *cd^g225^* heterozygous (knock-in,KI, blue), and *cd^225R^* homozygous (knock-out, KO, orange) mosquitoes. The survival after blood-feeding is represented by the Weibull parametric survival model with a single solid line per genotype (n = 271) and by the Kaplan-Meier survival curves (dashed line).

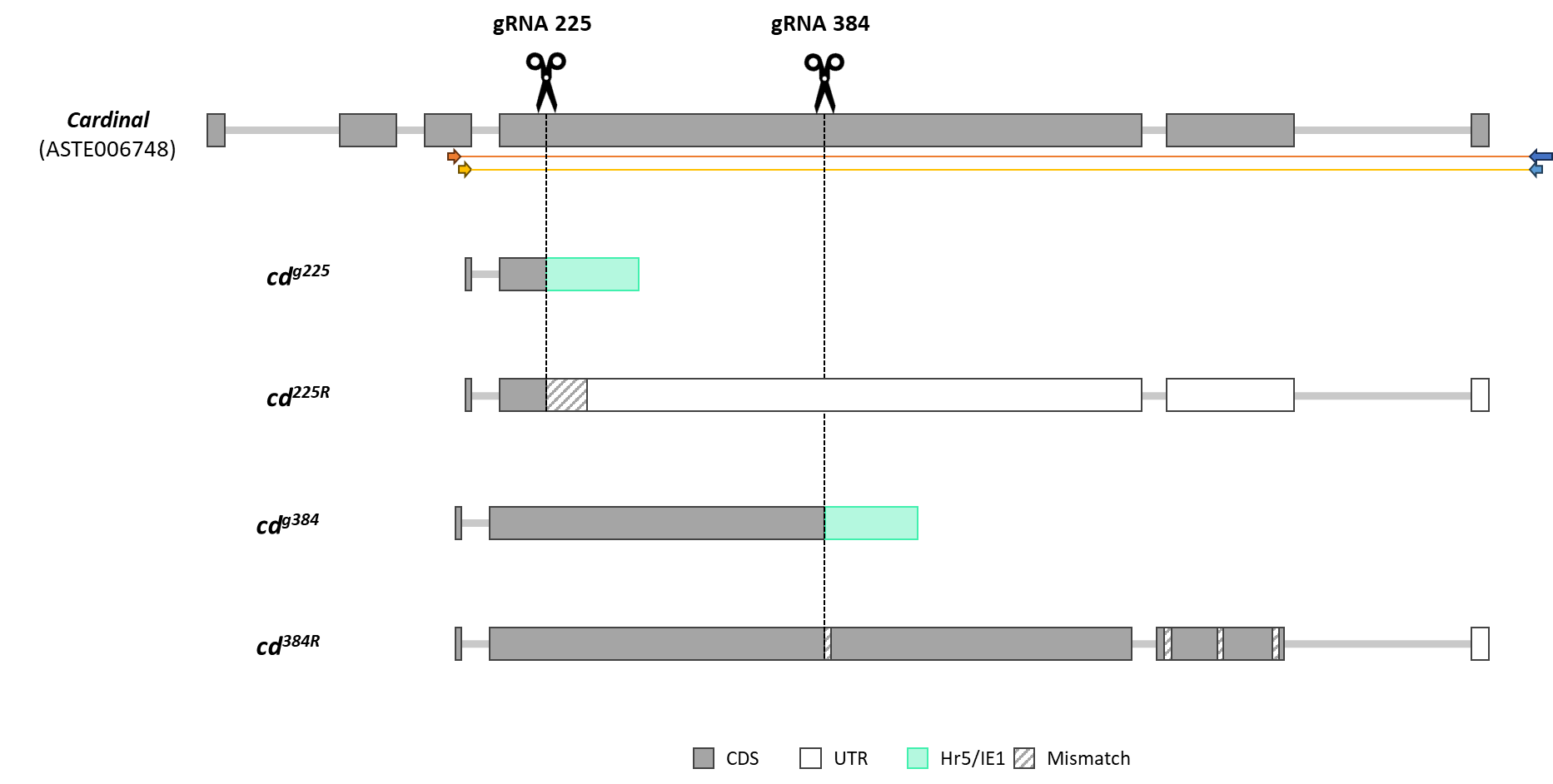

**SFig 3. Isoforms identified for the *cd^g225^, cd^g384^, cd^225R^, cd^384R^* mutant lines by a 3’ RACE PCR in comparison to the annotated ASTE006748 sequence.** The black scissors and the dashed line represent the target site of the two sgRNAs used to generate the four assessed lines. The orange arrow indicates the location of the GSP (LA8430), the yellow arrow indicates the alignment site of the ‘nested’ primer (LA8431), the long dark blue arrow represents the UPM, and the short light blue arrow represents the UPM short. The orange line represents the length of the expected amplicon for the 3’ RACE PCR and the yellow line represents the ‘nested’ PCR. The rectangular shape is indicative of exons while the connecting grey line represents the introns. The colour of the exons is different depending if they are a CDS (grey), an UTR (white) or part of the transgene (in this case Hr5/IE1, green). Translation of the obtained DNA sequence for the knock-out lines was aligned to the ASTE006748 annotated sequence to better characterise changes in the amino acid sequence. Regions that showed different amino acid sequences are represented in a diagonal grey pattern.

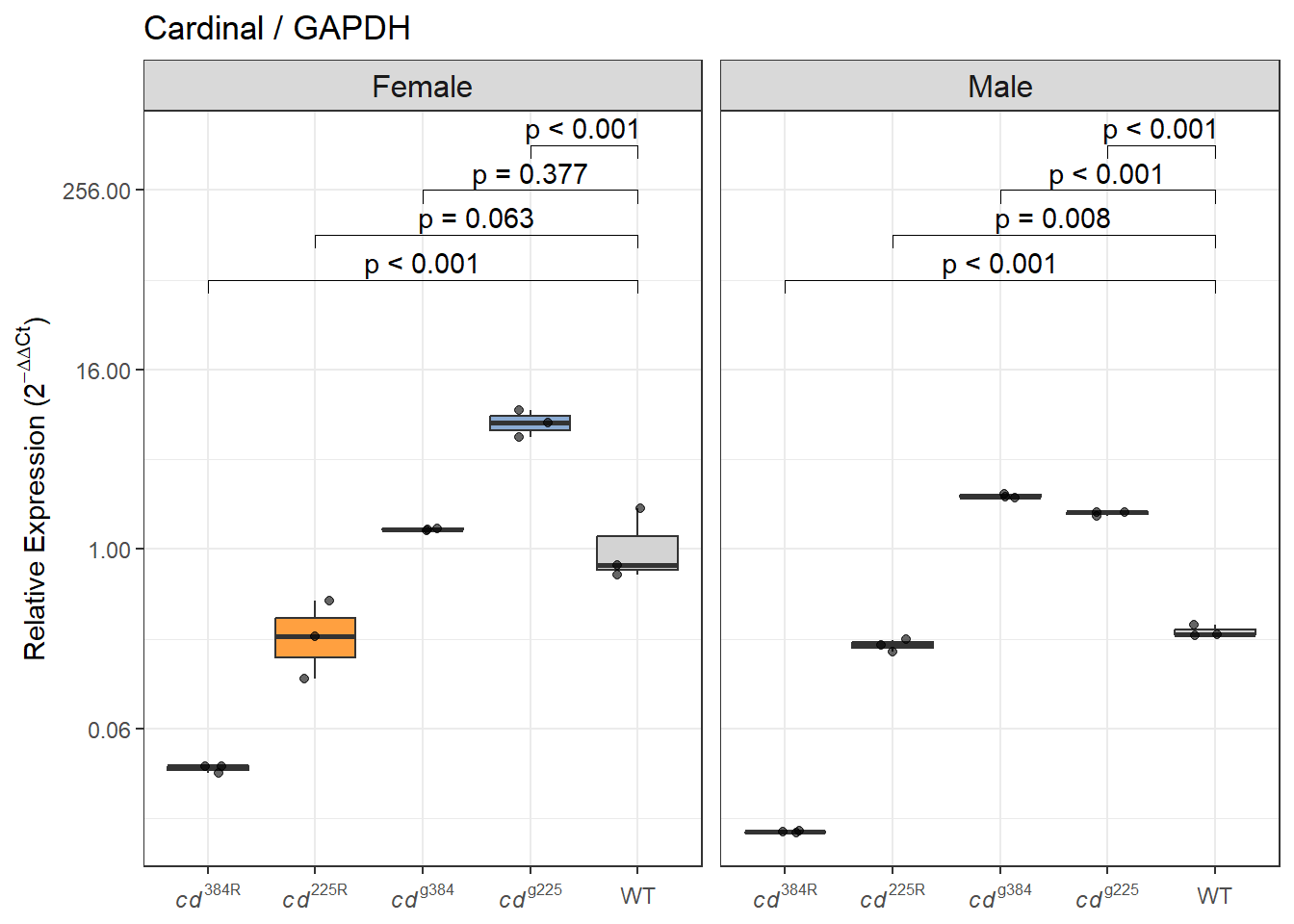

**SFig 4. The Hr5/IE1 promoter enhances *cd* expression in the knock-in lines.** Expression analysis of the *cd* gene by Taq-Man qPCR. Comparison of the expression levels of the *cd* gene of two different knock-in lines (*cd^g225^* and *cd^g384^*) to their respective knock-out lines (*cd^225R^* and *cd^384R^*) and using the WT female sample as a reference. *cd* expression was normalised against *GADPH* (housekeeping gene). The relative expression (RQ) is plotted on a log_2_-scaled axis. A permuted t-test with a 0.05 significance level was performed for each line in comparison to the control sample to determine statistical significance.

| *Predictors* | *Odds Ratios* | *CI* | *p* |
| --- | --- | --- | --- |
| Intercept  (*cd*^384R^ HET female) | 0.83 | 0.68 – 1.01 | 0.058 |
| Line  (*cd*^g384^ vs *cd*^384R^) | 1.22 | 0.93 – 1.61 | 0.157 |
| WT | 0.95 | 0.72 – 1.26 | 0.722 |
| HOM | 1.22 | 0.93 – 1.61 | 0.157 |
| Male | 1.15 | 0.87 – 1.52 | 0.321 |
| *cd*^g384^ × WT | 0.79 | 0.53 – 1.17 | 0.230 |
| *cd*^g384^ × HOM | 0.01 | 0.00 – 0.02 | **<0.001** |
| *cd*^g384^ × Male | 0.76 | 0.51 – 1.12 | 0.161 |
| WT × Male | 1.20 | 0.81 – 1.78 | 0.368 |
| HOM × Male | 0.71 | 0.48 – 1.05 | 0.089 |
| *cd*^g384^ × WT × Male | 1.42 | 0.81 – 2.48 | 0.215 |
| *cd*^g384^ × HOM × Male | 33.42 | 10.79 – 147.54 | **<0.001** |
| Observations | 24 | | |

**STable 1. Statistical analysis of larva to adult survival of *cd^g384^* and *cd^384R^* mosquitoes.** Homozygous viability was assessed by scoring adult eclosion. The proportion of adults eclosing out of an expected 200 was modelled with a binomial GLM including line (*cd^g384^* vs *cd^384R^*), genotype (HET, HOM, WT), sex, and all two- and three-way interactions (n = 24 observations). The reference group was *cd^384R^* heterozygous females.

|  | **Engorgement rate** | | | **Survival rate post blood-feeding** | | |
| --- | --- | --- | --- | --- | --- | --- |
| Cross (♀ X ♂) | *cd^384R^* | *cd^g384^* | *cd^g225^* | *cd^384R^* | *cd^g384^* | *cd^g225^* |
| SDA500 X SDA500 | 35/50 | n.d. | n.d. | 27/35 | n.d. | n.d. |
| SDA500 X WT† | n.d. | 43/50 | 22/50 | n.d. | 41/43 | 22/22 |
| SDA500 X Het | 39/50 | 50/50 | 39/50 | 32/39 | 48/50 | 39/39 |
| SDA500 X Hom | 29/50 | 41/50 | n.d. | 19/29 | 37/41 | n.d. |
| †WT X SDA500 | n.d. | 41/50 | 32/50 | n.d. | 34/41 | 32/32 |
| Het X SDA500 | 46/50 | 48/50 | 1/50 | 44/46 | 37/48 | 0/1 |
| Hom X SDA500 | 35/50 | 49/50 | n.d. | 32/35 | 27/49 | n.d. |

**STable 2. Numbers of females which took a blood meal (engorgement rate) and survived.** †wildtype siblings of transgenic lines; n.d. not determined.

A

| ***Predictors*** | ***Incidence Rate Ratios*** | ***CI*** | ***p*** |
| --- | --- | --- | --- |
| **Count Model** | | | |
| Intercept (*cd*^384R^ ,  SDA×SDA) | 76.59 | 62.78 – 93.45 | **<0.001** |
| *cd*^g225^ | 1.09 | 0.85 – 1.41 | 0.490 |
| *cd*^g384^ | 1.48 | 1.18 – 1.85 | **0.001** |
| Female Het | 1.31 | 1.03 – 1.67 | **0.029** |
| Male Het | 1.12 | 0.86 – 1.46 | 0.416 |
| Female Hom | 1.43 | 1.09 – 1.88 | **0.009** |
| Male Hom | 0.74 | 0.53 – 1.04 | 0.084 |
| *cd*^g225^ × Female Het | 1.00 | NaN – NaN | NaN |
| *cd*^g384^ × Female Het | 0.63 | 0.47 – 0.86 | **0.004** |
| *cd*^g225^ × Male Het | 0.90 | 0.62 – 1.31 | 0.569 |
| *cd*^g384^ × Male Het | 0.80 | 0.59 – 1.10 | 0.177 |
| *cd*^g384^ × Female Hom | 0.41 | 0.28 – 0.59 | **<0.001** |
| *cd*^g384^ × Male Hom | 0.79 | 0.50 – 1.25 | 0.316 |
| Intercept (*cd*^384R^, SDA×SDA) | 9534217.44 | 196910.85 – 1587058406.93 |  |
| **Zero-Inflated Model** | | | |
| Intercept (*cd*^384R^, SDA×SDA) | 0.47 | 0.39 – 0.56 | **<0.001** |
| **Random Effects** | | | |
| σ^2^ |  |  |  |
| τ_00_ _plate_well:rep_ | 0.05 | | |
| τ_00_ _rep_ | 0.00 | | |
| N _plate_well_ | 506 | | |
| N _rep_ | 10 | | |
| Observations | 546 | | |

**B**

| Line | Cross | **Non-zero wells** | | | Expected count (incl. zeros)*^1^* |
| --- | --- | --- | --- | --- | --- |
|  |  | Conditional mean | Lower 95% CI | Upper 95% CI |  |
| *cd*^384R^ | WT | 76.6 | 62.8 | 93.4 | 52.2 |
| *cd*^g225^ | WT | 83.7 | 71.6 | 97.9 | 57.0 |
| *cd*^g384^ | WT | 113.4 | 102.2 | 125.9 | 77.3 |
| *cd*^384R^ | Female HET | 100.5 | 87.1 | 115.8 | 68.4 |
| *cd*^g225^ | Female HET | 109.7 | NaN | NaN | 74.8 |
| *cd*^g384^ | Female HET | 94.4 | 80.1 | 111.2 | 64.3 |
| *cd*^384R^ | Male HET | 85.5 | 71.6 | 102.2 | 58.3 |
| *cd*^g225^ | Male HET | 83.8 | 67.5 | 104.0 | 57.1 |
| *cd*^g384^ | Male HET | 101.8 | 88.4 | 117.2 | 69.4 |
| *cd*^384R^ | Female HOM | 109.9 | 91.1 | 132.4 | 74.8 |
| *cd*^g225^ | Female HOM | NA | NA | NA | NA |
| *cd*^g384^ | Female HOM | 66.5 | 53.2 | 83.2 | 45.3 |
| *cd*^384R^ | Male HOM | 56.9 | 43.4 | 74.7 | 38.8 |
| *cd*^g225^ | Male HOM | NA | NA | NA | NA |
| *cd*^g384^ | Male HOM | 66.7 | 49.9 | 89.3 | 45.5 |
| *^1^* Structural zero probability = 0.319; expected count = conditional mean × (1 − p_zero) | | | | | |

| **C Line comparisons within cross** | | | | | | |
| --- | --- | --- | --- | --- | --- | --- |
| Contrast | Cross | IRR | Lower 95% CI | Upper 95% CI | z | p |
| *cd*^384R^ - *cd*^g225^ | WT | 0.91 | 0.71 | 1.18 | −0.69 | 0.769 |
| *cd*^384R^ - *cd*^g384^ | WT | 0.68 | 0.54 | 0.84 | −3.45 | 0.002 |
| *cd*^g225^ - *cd*^g384^ | WT | 0.74 | 0.61 | 0.89 | −3.19 | 0.004 |
| *cd*^384R^ - *cd*^g384^ | Female HET | 1.06 | 0.86 | 1.32 | 0.57 | 0.568 |
| *cd*^384R^ - *cd*^g225^ | Male HET | 1.02 | 0.77 | 1.35 | 0.14 | 0.989 |
| *cd*^384R^ - *cd*^g384^ | Male HET | 0.84 | 0.67 | 1.05 | −1.52 | 0.283 |
| *cd*^g225^ - *cd*^g384^ | Male HET | 0.82 | 0.64 | 1.06 | −1.49 | 0.297 |
| *cd*^384R^ - *cd*^g384^ | Female HOM | 1.65 | 1.24 | 2.21 | 3.40 | <0.001 |
| *cd*^384R^ - *cd*^g384^ | Male HOM | 0.85 | 0.57 | 1.27 | −0.78 | 0.433 |

| **D Cross comparisons within line** | | | | | | |
| --- | --- | --- | --- | --- | --- | --- |
| Contrast | Line | IRR | Lower 95% CI | Upper 95% CI | z | p |
| WT - Female HET | *cd*^384R^ | 0.76 | 0.60 | 0.97 | −2.19 | 0.184 |
| WT - Male HET | *cd*^384R^ | 0.90 | 0.69 | 1.17 | −0.81 | 0.927 |
| WT - Female HOM | *cd*^384R^ | 0.70 | 0.53 | 0.91 | −2.60 | 0.070 |
| WT - Male HOM | *cd*^384R^ | 1.35 | 0.96 | 1.88 | 1.73 | 0.415 |
| Female HET - Male HET | *cd*^384R^ | 1.17 | 0.94 | 1.47 | 1.40 | 0.630 |
| Female HET - Female HOM | *cd*^384R^ | 0.91 | 0.72 | 1.15 | −0.75 | 0.943 |
| Female HET - Male HOM | *cd*^384R^ | 1.77 | 1.30 | 2.40 | 3.63 | 0.003 |
| Male HET - Female HOM | *cd*^384R^ | 0.78 | 0.60 | 1.01 | −1.91 | 0.310 |
| Male HET - Male HOM | *cd*^384R^ | 1.50 | 1.09 | 2.08 | 2.46 | 0.100 |
| Female HOM - Male HOM | *cd*^384R^ | 1.93 | 1.39 | 2.68 | 3.91 | <0.001 |
| WT - Male HET | *cd*^g225^ | 1.00 | 0.77 | 1.30 | −0.01 | 0.994 |
| WT - Female HET | *cd*^g384^ | 1.20 | 0.99 | 1.45 | 1.88 | 0.325 |
| WT - Male HET | *cd*^g384^ | 1.11 | 0.94 | 1.32 | 1.23 | 0.737 |
| WT - Female HOM | *cd*^g384^ | 1.70 | 1.34 | 2.18 | 4.28 | <0.001 |
| WT - Male HOM | *cd*^g384^ | 1.70 | 1.25 | 2.31 | 3.37 | 0.007 |
| Female HET - Male HET | *cd*^g384^ | 0.93 | 0.75 | 1.15 | −0.69 | 0.958 |
| Female HET - Female HOM | *cd*^g384^ | 1.42 | 1.08 | 1.87 | 2.49 | 0.092 |
| Female HET - Male HOM | *cd*^g384^ | 1.41 | 1.01 | 1.97 | 2.04 | 0.248 |
| Male HET - Female HOM | *cd*^g384^ | 1.53 | 1.18 | 1.99 | 3.18 | 0.013 |
| Male HET - Male HOM | *cd*^g384^ | 1.53 | 1.10 | 2.11 | 2.56 | 0.077 |
| Female HOM - Male HOM | *cd*^g384^ | 1.00 | 0.69 | 1.44 | −0.02 | 1.000 |

**STable 3. Statistical analysis of fecundity data.** Fecundity (egg count per well) was modelled using a zero-inflated negative binomial mixed model with cross direction and genetic line as fixed effects, and replicate and plate well as nested random effects (n = 341 observations across 340 wells, 10 replicates). Model derived estimated means with confidence intervals (B) and post hoc comparisons follow (C, D). NaN: Not a number, confidence intervals inestimable for categories with no eggs.

| **A**  *Predictors* | *Odds Ratios* | *CI* | *p* |
| --- | --- | --- | --- |
| Intercept (*cd*^384R^,  SDA×SDA) | 4.21 | 2.16 – 8.22 | **<0.001** |
| *cd*^g225^ | 0.88 | 0.37 – 2.09 | 0.772 |
| *cd*^g384^ | 1.28 | 0.59 – 2.77 | 0.539 |
| Female Het | 4.51 | 1.91 – 10.65 | **0.001** |
| Male Het | 2.02 | 0.81 – 5.07 | 0.132 |
| Female Hom | 2.96 | 1.11 – 7.88 | **0.030** |
| Male Hom | 1.92 | 0.66 – 5.61 | 0.230 |
| *cd*^g384^ × Female Het | 0.15 | 0.05 – 0.45 | **0.001** |
| *cd*^g225^ × Male HET | 0.37 | 0.10 – 1.35 | 0.131 |
| *cd*^g384^ × Male Het | 0.31 | 0.10 – 0.96 | **0.043** |
| *cd*^g384^ × Female Hom | 0.00 | 0.00 – 0.01 | **<0.001** |
| *cd*^g384^ × Male Hom | 0.30 | 0.07 – 1.31 | 0.108 |
| **Random Effects** | | | |
| σ^2^ | 2.76 | | |
| τ_00_ _plate_well:rep:line_ | 2.72 | | |
| τ_00_ _rep:line_ | 0.00 | | |
| τ_00_ _line_ | 0.00 | | |
| N _plate_well_ | 424 | | |
| N _rep_ | 10 | | |
| N _line_ | 3 | | |
| Observations | 449 | | |

**B**

| Line | Cross | **Estimated marginal means** | | |
| --- | --- | --- | --- | --- |
|  |  | Hatching probability | Lower 95% CI | Upper 95% CI |
| *cd*^384R^ | WT | 0.808 | 0.683 | 0.892 |
| *cd*^g225^ | WT | 0.787 | 0.681 | 0.865 |
| *cd*^g384^ | WT | 0.843 | 0.784 | 0.888 |
| *cd*^384R^ | Female HET | 0.950 | 0.917 | 0.970 |
| *cd*^g225^ | Female HET | NA | NA | NA |
| *cd*^g384^ | Female HET | 0.781 | 0.665 | 0.865 |
| *cd*^384R^ | Male HET | 0.895 | 0.820 | 0.941 |
| *cd*^g225^ | Male HET | 0.733 | 0.563 | 0.853 |
| *cd*^g384^ | Male HET | 0.773 | 0.671 | 0.851 |
| *cd*^384R^ | Female HOM | 0.926 | 0.859 | 0.962 |
| *cd*^g225^ | Female HOM | NA | NA | NA |
| *cd*^g384^ | Female HOM | 0.048 | 0.022 | 0.101 |
| *cd*^384R^ | Male HOM | 0.890 | 0.779 | 0.949 |
| *cd*^g225^ | Male HOM | NA | NA | NA |
| *cd*^g384^ | Male HOM | 0.754 | 0.541 | 0.888 |

| **C Line comparisons within cross** | | | | | | |
| --- | --- | --- | --- | --- | --- | --- |
| Contrast | Cross | OR | Lower  95% CI | Upper  95% CI | z | p |
| *cd*^384R^ - *cd*^g225^ | WT | 1.14 | 0.48 | 2.70 | 0.29 | 0.955 |
| *cd*^384R^ - *cd*^g384^ | WT | 0.78 | 0.36 | 1.70 | −0.62 | 0.812 |
| *cd*^g225^ - *cd*^g384^ | WT | 0.69 | 0.35 | 1.35 | −1.08 | 0.527 |
| *cd*^384R^ - *cd*^g384^ | Female HET | 5.33 | 2.41 | 11.81 | 4.13 | <0.001 |
| *cd*^384R^ - *cd*^g225^ | Male HET | 3.11 | 1.17 | 8.30 | 2.27 | 0.060 |
| *cd*^384R^ - *cd*^g384^ | Male HET | 2.50 | 1.11 | 5.64 | 2.21 | 0.070 |
| *cd*^g225^ - *cd*^g384^ | Male HET | 0.80 | 0.32 | 2.00 | −0.47 | 0.885 |
| *cd*^384R^ - *cd*^g384^ | Female HOM | 246.12 | 84.60 | 716.03 | 10.11 | <0.001 |
| *cd*^384R^ - *cd*^g384^ | Male HOM | 2.65 | 0.75 | 9.41 | 1.51 | 0.132 |

| **D Cross comparisons within line** | | | | | | |
| --- | --- | --- | --- | --- | --- | --- |
| Contrast | Line | OR | Lower 95% CI | Upper  95% CI | z | p |
| WT - Female HET | *cd*^384R^ | 0.22 | 0.09 | 0.52 | −3.43 | 0.005 |
| WT - Male HET | *cd*^384R^ | 0.49 | 0.20 | 1.24 | −1.51 | 0.559 |
| WT - Female HOM | *cd*^384R^ | 0.34 | 0.13 | 0.90 | −2.17 | 0.190 |
| WT - Male HOM | *cd*^384R^ | 0.52 | 0.18 | 1.51 | −1.20 | 0.752 |
| Female HET - Male HET | *cd*^384R^ | 2.23 | 0.97 | 5.10 | 1.89 | 0.320 |
| Female HET - Female HOM | *cd*^384R^ | 1.52 | 0.62 | 3.73 | 0.92 | 0.889 |
| Female HET - Male HOM | *cd*^384R^ | 2.34 | 0.87 | 6.33 | 1.68 | 0.448 |
| Male HET - Female HOM | *cd*^384R^ | 0.68 | 0.26 | 1.77 | −0.78 | 0.936 |
| Male HET - Male HOM | *cd*^384R^ | 1.05 | 0.37 | 2.99 | 0.09 | 1.000 |
| Female HOM - Male HOM | *cd*^384R^ | 1.54 | 0.51 | 4.62 | 0.77 | 0.940 |
| WT - Male HET | *cd*^g225^ | 1.35 | 0.53 | 3.44 | 0.63 | 0.526 |
| WT - Female HET | *cd*^g384^ | 1.51 | 0.75 | 3.04 | 1.15 | 0.781 |
| WT - Male HET | *cd*^g384^ | 1.57 | 0.82 | 3.01 | 1.37 | 0.645 |
| WT - Female HOM | *cd*^g384^ | 106.02 | 43.85 | 256.32 | 10.35 | <0.001 |
| WT - Male HOM | *cd*^g384^ | 1.76 | 0.63 | 4.92 | 1.07 | 0.822 |
| Female HET - Male HET | *cd*^g384^ | 1.04 | 0.48 | 2.28 | 0.11 | 1.000 |
| Female HET - Female HOM | *cd*^g384^ | 70.29 | 26.25 | 188.21 | 8.46 | <0.001 |
| Female HET - Male HOM | *cd*^g384^ | 1.16 | 0.38 | 3.56 | 0.27 | 0.999 |
| Male HET - Female HOM | *cd*^g384^ | 67.36 | 26.17 | 173.41 | 8.73 | <0.001 |
| Male HET - Male HOM | *cd*^g384^ | 1.12 | 0.38 | 3.30 | 0.20 | 1.000 |
| Female HOM - Male HOM | *cd*^g384^ | 0.02 | 0.00 | 0.06 | −6.49 | <0.001 |

**STable 4. Statistical analysis of fertility assay.** Hatching rate (larvae per egg) was modelled using a binomial GLMM with a combined line × cross factor as the fixed effect, and nested random effects of plate well within replicate within line-cross group. All comparisons are relative to WT (A). Model estimated means with confidence intervals (B) and post hoc comparisons (C, D).

| *Predictors* | *Estimates* | *CI* | *p* |
| --- | --- | --- | --- |
| shape | 1.38 | 0.73 – 2.60 | 0.327 |
| scale | 1127.37 | 0.00 – 1.5x10^230^ | 0.979 |
| *cd*^g384^ Het | 1.49 | 0.93 – 2.36 | 0.095 |
| *cd*^g384^ Hom | 0.40 | 0.25 – 0.63 | **<0.001** |
| *cd*^g384^ WT | 2.03 | 1.28 – 3.23 | **0.003** |
| *cd*^g384del^ Het | 0.20 | 0.12 – 0.32 | **<0.001** |
| *cd*^g384del^ WT | 1.19 | 0.75 – 1.90 | 0.457 |
| *cd*^g338-384^ Het | 0.07 | 0.04 – 0.11 | **<0.001** |
| *cd*^g338-384^ WT | 514758.94 | 0.00 – Inf | 0.996 |
| *cd*^g225^ Het | 0.09 | 0.06 – 0.14 | **<0.001** |
| *cd*^g225^ WT | 2.73 | 1.71 – 4.34 | **<0.001** |
| *cd*^225R^ WT | 0.70 | 0.44 – 1.11 | 0.130 |
| Observations | 384 | | |

**STable 5. Statistical analysis for post-blood-feeding survival.** Post-blood-feeding survival was modelled using a Weibull parametric survival model with a single combined line × genotype factor (n = 384). Model estimates are scale factors relative to the reference group (*cd^225R^* Hom); values below 1 indicate shorter survival relative to this reference.

| *Predictors* | *Odds Ratios* | *CI* | *p* |
| --- | --- | --- | --- |
| Intercept (WT) | 0.06 | 0.02 – 0.13 | **<0.001** |
| KI | 65.61 | 25.70 – 205.20 | **<0.001** |
| KO | 0.00 | NA – 3.179x10^175^ | 0.996 |
| Observations | 9 | | |

**STable 6. Statistical analysis of Smurf assay.** The proportion of smurf-positive mosquitoes (indicating gut barrier disruption) was modelled using a binomial GLM with genotype (WT, KI, KO) as the sole predictor, restricted to sucrose-delivered mosquitoes (n = 9 cage-level observations). The reference group was WT.

| **A**  *Predictors* | *Estimates* | *CI* | *p* |
| --- | --- | --- | --- |
| shape | 2.17 | 1.27 – 3.71 | **0.004** |
| scale | 95.42 | 0.00 – 356311913167.43 | 0.685 |
| Sugar supplementation | 1.65 | 1.27 – 2.15 | **<0.001** |
| 6 mM NaOH control | 0.9 | 0.68 – 1.19 | 0.468 |
| 6 mM XA | 1.64 | 1.15 – 2.34 | **0.007** |
| WT | 5.17 | 1.91 – 14.01 | **<0.001** |
| Observations | 211 | | |

| **B**  Experiment | Dose | Genotype | **Survival probability at 72 h** | | |
| --- | --- | --- | --- | --- | --- |
|  |  |  | Survival | Lower 95% CI | Upper 95% CI |
| Blood supplementation | | | | | |
| blood supplementation | 0mM | Het | 0.582 | 0.407 | 0.733 |
| blood supplementation | 6mM | Het | 0.831 | 0.701 | 0.907 |
| blood supplementation | 6mM C- | Het | 0.507 | 0.336 | 0.677 |
| blood supplementation | 0mM | WT | 0.985 | 0.884 | 0.998 |
| blood supplementation | 6mM | WT | 0.995 | 0.963 | 0.999 |
| Sugar supplementation | | | | | |
| sugar supplementation | 0mM | Het | 0.834 | 0.727 | 0.906 |
| sugar supplementation | 6mM | Het | 0.940 | 0.878 | 0.972 |
| sugar supplementation | 6mM C- | Het | 0.796 | 0.682 | 0.879 |
| sugar supplementation | 0mM | WT | 0.995 | 0.965 | 0.999 |
| sugar supplementation | 6mM | WT | 0.998 | 0.987 | 1.000 |

**STable 7. Statistical analysis of survival after XA supplementation.** The effect of xanthurenic acid (XA) supplementation on post-blood-feeding survival was examined using a Weibull parametric survival model (n = 387). Predictors were supplementation route (blood vs sugar), XA dose (0 mM, 6 mM NaOH control [6mM C−], or 6 mM XA), and genotype (Het vs WT). The reference group was Het mosquitoes receiving blood supplementation at 0 mM (A). Model predicted survival probability at 72h, with confidence intervals (B)

| *Predictors* | *Estimates* | *CI* | *p* |
| --- | --- | --- | --- |
| shape | 1.335 | 0.411 – 4.334 | 0.630 |
| scale | 70.891 | 0.000 – 9.8x10^28^ | 0.894 |
| *cd*^g225^ WT | 26.664 | 11.043 – 64.385 | **<0.001** |
| Trans-het (*cd*^225R^:*cd*^g225^ Het) | 0.969 | 0.401 – 2.340 | 0.944 |
| Het^KO^ (*cd*^225R^:*cd*^g225^ WT) | 12218566.257 | 0.000 – Inf | 0.997 |
| Observations | 186 | | |

**STable 8. Statistical analysis of survival.** To assess whether the post-blood-feeding survival phenotype is caused by disruption of cardinal function, survival data were modelled for the *cd^g225^* and *cd^225R^* lines specifically, using a Weibull parametric model (n = 186). Predictors were line (*cd^g225^* vs *cd^225R^*) and genotype (Het, Hom, WT), with the reference group being *cd^g225^* heterozygotes.
